# MicroRNA-502-3p modulates the GABA A subunits, synaptic proteins and mitochondrial morphology in hippocampal neurons

**DOI:** 10.1101/2025.01.09.632229

**Authors:** Bhupender Sharma, Daniela Rodarte, Gunjan Goyal, Sheryl Rodriguez, Subodh Kumar

## Abstract

MicroRNA-502-3p (MiR-502-3p), a synapse enriched miRNA is considerably implicated in Alzheimer’s disease (AD). Our previous study found the high expression level of miR-502-3p in AD synapses relative to controls. Further, miR-502-3p was found to modulate the GABAergic synapse function via modulating the GABA A receptor subunit α-1 (GABRA1) protein. The current study is attempted to examine the impact of miR-502-3p on other GABA subunit proteins, synaptic proteins, mitochondrial morphology and other hippocampal neuron genes. Mouse hippocampal neuronal (HT22) cells were transfected with miR-502-3p overexpression (OE) vector, miR-502-3p sponge (suppression) vector and scramble control vector. MiR-502-3p vectors transfection was confirmed by fluorescence microscopy. MiR-502-3p expression and *GABRA1* expression was confirmed by qRT-PCR and miRNAScope *in-situ* hybridization. GABA A subunit and synaptic proteins were studied by immunoblotting analysis and mitochondrial morphology was analyzed by transmission electron microscopy (TEM) analysis. Further, Affymetrix gene array analysis was conducted in miR-502-3p overexpressed and suppressed cells. Our results observed that elevated miR-502-3p, negatively modulates the GABRA1 level. The levels of GABA A subunit and synaptic proteins were reduced by ectopic expression of miR-502-3p and increase by miR-502-3p suppression. The mitochondrial morphology was found to be improved in-terms of their number and length in miR-502-3p suppressed cells. Further, Gene array analysis unveiled the deregulation of several genes by miR-502-3p, which are associated with oxidative stress, immune response and synaptic function. These results provide new insights and an update to understand the biological roles of miR-502-3p in regulation of neuron function and synaptic activity.

## INTRODUCTION

MicroRNAs (miRNAs) are small (∼22 nucleotides), non-coding single-stranded RNAs that regulate gene expression at the post-transcriptional level. They bind to the 3’ untranslated regions (UTRs) of target messenger RNAs (mRNAs), leading to mRNA degradation or translational repression. Dysregulation of miRNAs has been extensively reported in several diseases along with neurodegenerative diseases, including Alzheimer’s disease (AD) (1,2). Changes in miRNA expression can affect processes such as amyloid-beta (Aβ) metabolism, tau phosphorylation, and synaptic function, contributing to the pathogenesis of AD (3,4). Synaptic dysfunction plays a critical role in the early stages of AD, and the impairment of inhibitory neurotransmission mediated by the gamma-aminobutyric acid (GABA) system has been increasingly implicated in this process (5,6). Synaptic dysfunction is characterized by a loss of synaptic connections and alterations in synaptic plasticity (7,8). Synaptic dysfunction is closely linked to cognitive decline and is often observed before significant neuronal loss (9). Changes in synaptic structure and function are thought to result from the toxic effects of Aβ plaques (10–12) and tau tangles, as well as disruptions in neurotransmitter systems (13–15).

Our lab recently explored the role of microRNA-502-3p (miR-502-3p) in AD pathogenesis (16–19). In previous research we found overexpressed miR-502-3p/miR-501-3p levels in AD postmortem brain synaptosomes as compared to unaffected controls. We observed that miR-502-3p and miR-501-3p showed positive correlation with AD severity in term of Braak stages-based disease progression (16). By *in-silico* analysis we found that miR-500 family targets GABA function, neural function and synaptic functions (16,19). We have chosen miR-502-3p for further investigation because it is more conserved in human and mouse and closely associated to brain function as confirmed by Ingenuity pathway analysis (16,18). Moreover, miR-502-3p is least investigated in AD and synaptic function. GABAergic neurons, the most prevalent inhibitory neurons in the human brain, experience reduction in various neurological disorders, including AD and AD-related dementia (ADRD) (5). Our recent research found that miR-502-3p may contribute to the pathogenesis of AD by regulating the expression of gamma-aminobutyric acid type A receptor subunit α-1 (GABRA1), which is crucial for inhibitory neurotransmission in the brain. GABRA1 is a primary component of GABA A receptors, the major inhibitory neurotransmitter receptors in the central nervous system. These receptors mediate synaptic inhibition by facilitating chloride ion influx, hyperpolarizing neurons and reducing their excitability. Our research evidence indicates that miR-502-3p can bind to the 3’ UTR of GABRA1 mRNA, downregulating its expression and potentially contributing to synaptic dysfunction in AD. Moreover, miR-502-3p modulates the hippocampal neuron survival and positively modulates the GABA current (19).

Since miR-502-3p is a synapse localized miRNA, hence more research is needed to investigate the role of miR-502-3p in synaptic dysfunction in AD. The current study is an extension of ongoing research project in our lab with focus on miR-502-3p and synaptic function. Our study found some interesting findings when we modulated the miR-502-3p level (overexpression and suppression) in the hippocampal neurons. MiR-502-3p was found to be implicated in modulation of GABA A subunit proteins, synaptic proteins, mitochondrial morphology and gene expression changes. These finding will aid to understand the underlying molecular mechanism of miR-502-3p at synapse and unveil its therapeutic potential in AD and neurodegenerative diseases.

## MATERIALS AND METHODS

### Cell Culture

The mouse hippocampal (HT22) neuronal cells were routinely maintained in our lab (19). Cells were cultured in Dulbecco’s modified eagle medium (DMEM, Thermo Fisher Scientific) supplemented with 10% fetal bovine serum (FBS, Gibco) and 1% penicillin/ streptomycin (complete media) at 37^°^C in a humidified atmosphere with 5% CO₂. HT22 cells (0.2 x 10^6^ cells/ well) were seeded in a 6-well plate with DMEM supplemented with 5% FBS without antibiotics. After 8 hours, when the cells had adhered to the surface, they were transfected using lipofectamine 3000 (Invitrogen) according to the manufacturer’s protocol (19).

### MiR-502-3p overexpression (OE) and miR-502-3p suppression (sponge) vector

The HT22 cells were categorized into three groups as they were transfected with three different plasmids (VectorBuilder, USA). These included (i) scramble control plasmid (pRP[Exp]- CAG>EGFP) with GFP expression, (ii) miR-502-3p overexpression (OE) plasmid (pRP[shRNA]- EGFP-U6>{hsa-microRNA-502-3p}) for overexpression of hsa-miR-502-3p along with GFP expression, and (iii) sponge plasmid (pRP[Exp]-CAG>EGFP:miRNA-502-3p sponge) designed to inhibit hsa-miR-502-3p by containing multiple binding sites for this miRNA, also with GFP expression. In this study we used miR-502-3p expression/ inhibition plasmids engineered with fluorescent markers and inducible promoters, providing more stable and controlled expression of the miR-502-3p. Complete details of miR-502-3p vectors are provided in supplementary information (**Supplementary information file 1**). This stability ensures consistent effects on cellular processes, gene expression, and phenotype over extended periods. Following 24 hours of post-transfection, complete DMEM media was added, and the transfected HT22 cells were cultured for 48 h at 37^°^C in a humidified atmosphere with 5% CO_₂_. To assess the transfection efficiency, GFP expression was examined 48 hours post-transfection using fluorescence microscopy (AMG Evos Fl Microscope, E1512-155E-088). GFP-positive cells were quantified using ImageJ software to determine transfection efficiency.

### qRT-PCR analysis

Quantitative reverse transcription PCR (qRT-PCR) was performed to quantify the expression of miR-502-3p in transfected HT22 cells. Total RNA was extracted from HT22 cells using TriZol reagent (Invitrogen, USA) and quantified by nanodrop. Thereafter, for miRNA analysis, 2 ug of RNA was polyadenylated and reverse transcribed into cDNA using the miRNA first-strand synthesis kit (Agilent) according to manufacturer’s protocol (19). For GABRA1 expression analysis, cDNA was synthesized from 2 ug of RNA by using SuperScript III first – strand synthesis kit (Invitrogen). The miRNAs and gene specific primers were commercially synthesized by Integrated DNA Technologies (IDT), Coralville, IA, USA, **Supplementary information file 2**). The qRT-PCR was conducted by preparing a reaction mixture that included 1 μL of miRNA / or GABRA1 gene specific forward primer (10 μM; IDT), 1 μL of universal reverse primer (3.125 μM; Agilent) or GABRA1 gene specific reverse primer (10 μM; IDT), 5 μL of 2 × SYBR Green PCR Master Mix (KAPA SYBR, Roche), and 1 μL of cDNA. RNase-free water was added to bring the final volume to 10 μL. U6 (SnRNA) and Beta actin expression was also measured in the cells to normalize miRNA and mRNA expression as an internal control. Each sample was prepared in triplicate and analyzed using the LightCycler 96 Real-Time PCR (Roche Diagnostics, Indianapolis, IN, USA). The qRT-PCR protocol for miRNA and GABRA1 gene expression involved an initial denaturation at 95^°^C for 10 minutes, followed by 50 cycles of 95^°^C for 10 seconds, 60^°^C for 20 seconds, and 72^°^C for 20 seconds. The miRNA/ mRNA fold change was calculated using the formula (2^^−ΔΔct^) (19).

### MicroRNA *in-situ* hybridization

To assess the co-localization of miR-502-3p and GABRA1 within transfected HT22 cells, we performed miRNA *in-situ* Hybridization using RNAScope™ (Advanced Cell Diagnostics, Newark, CA, USA) (19). HT22 cells were seeded onto polyl-lysine coated glass coverslips at a density of approximately 0.1 × 10^5^ cells/ well, then transfected with scramble control, miR-502-3p overexpression and miR-502-3p sponge plasmids using lipofectamine. Forty-two hours post transfection, cells were fixed with 10% neutral buffered formalin, permeabilized with PBS containing 0.1% Tween 20, and hybridized with target probes for hsa-miR-502-3p, mmu-U6, or a scramble control. After hybridization, signal amplification was carried out as per the manufacturer’s instructions (Advanced Cell Diagnostics; Newark, CA, USA). For GABRA1 detection cells fixed on coverslip were incubated overnight with GABRA1 primary antibody and HRP-conjugated secondary antibody anti-rabbit-HRP. The details of antibodies and their dilutions used are listed in **Supplementary information file 2**. After washing the cells five times with TBS-T buffer at 10-minute intervals, cells were detected with 2.5 LS Duplex green B (Advanced Cell Diagnostics, Newark, CA, USA). Thereafter the samples were then stained with 50% hematoxylin, mounted with VectaMount mounting medium (Vector Laboratories, Newark, CA, USA), and visualized using a fluorescent motorized upright microscope (Leica microsystem).

### Immunoblotting analysis

Transfected HT22 cells pellet(s) were collected by trypsinization and washed with PBS. Thereafter HT22 cell pellet(s) were suspended in RIPA buffer (Thermo Scientific) supplemented with protease inhibitors (Thermo Scientific) and disrupted by ultra-sonication (Qsonica, USA; amplitude 80%, pulse 10 sec on/ off, time 10 seconds). Thereafter, cell debris were removed by centrifugation (10,000 rpm for 10 minutes). Protein concentration was estimated by BCA assay (Thermo Scientific). Equal amounts of protein (40 μg per sample) were separated by SDS-PAGE on 10% polyacrylamide gels and transferred onto PVDF membranes (BioRad, Hercules, CA, USA). Membranes were blocked in 5% BSA for 1 hour at room temperature. The membranes were then incubated overnight at 4^°^C with primary antibodies against GABRA1, GABRB3, GABRG2, Calmodulin, Gephryin, SNAP25, Syntaxin1, MAP2, NRXN1 and VAMP2 (1:1000 v/v, Proteintech). Details of the antibodies and their dilutions used are listed in **Supplementary information file 2**. After washing the membranes three times with TBS-T buffer at 10-minute intervals, they were incubated for 1 hour at room temperature with a secondary antibody (rabbit anti-mouse horseradish peroxidase [HRP] 1:10,000). Following three additional washes with TBS-T buffer, proteins were detected using chemiluminescence reagents (Thermo Fisher Scientific, USA) and visualized with an Amersham imager 680 (GE Healthcare Bio-Sciences, Uppsala, Sweden). Protein band intensities were quantified using ImageJ software (1.54d, Java 1.8.0_345: http://imagej.org) for densitometry analysis as described in more detail in our earlier study (19), and relative protein expression levels were normalized to β-actin as loading controls.

### Transmission Electron Microscopy

Electron microscopy experiments were performed on HT22 cells in 60-mm Petri dishes. Cells were transfected with scramble control vector, miR-502-3p overexpression vector and miR-502-3p suppression sponge vector. At 48 hours post-transfection, the cells were washed with 5 mL 1X phosphate buffer saline. Cells were scraped out in a fixative solution (8% glutaraldehyde, 16% paraformaldehyde, and 0.2M sodium cacodylate buffer) and pellet was dissolved in fixative solution for 1 h at room temperature. The cells were centrifuged, and pellet were dissolved into fresh fixative solution. Cells were incubated at room temperature for 30 min following centrifugation at 300 g for 3 min (20). The resulting cell pellet was processed for electron microscopy analysis for mitochondria structure and morphology at the Imaging Core Facility at Texas Tech University, Lubbock, TX, USA.

### Affymetrix Gene Array analysis

Total RNA was extracted from transfected HT22 cells (scramble control, miR-502-3p overexpression, and miR-502-3p sponge plasmids) using the TriZol reagent (Invitrogen, USA). The RNA quantity and purity was determined by NanoDrop assay. The RNA samples were processed for Affymetrix gene array analysis using Genomic and microarray core facility at UT Southwestern Medical Center, Dallas TX (www.microarray.swmed.edu). Clariom S assay mouse platform was used to access the gene expression changes in mouse hippocampal neuron cells treated with scramble control, miR-502-3p OE and miR-502-3p sponge vector in duplicates. Clariom S assays focus on well-annotated genes (>20,000), providing researchers with the ability to perform gene-level expression profiling studies and to quickly assess changes in key genes and pathways.

### Gene array data analysis

Raw data were obtained, using the Affymetrix GeneChip array in the form of an individual CEL files. Each sample was then analyzed, using Transcriptome Analysis Console software v. 4. Tukey’s bi-weight average (log2) intensity was analyzed with an ANOVA P-value (<0.05) and FDR P-value (<0.05) for all conditions, for all genes in the samples from scramble control, miR-502-3p OE vector and miR-502-3p sponge vector treated group. SAM (significance analysis of microarray) with the R package was used to identify differentially expressed mRNA and gene probe sets in samples from the control, OE and sponge groups. Probe sets were considered biologically significant if the fold changes were >+2 and <-2. A Probe set (Gene/Exon) is considered expressed if ≥ <u>50</u>% samples have DABG values below DABG Threshold. DABG < 0.05 is considered statistically significant (21).

### Bioinformatics analysis of gene array data

Gene ontology enrichment analysis is conducted for deregulated genes by using the ShinyGO 0.81 software (16). The species category was selected for mouse. The deregulated genes obtained in three different comparisons (miR-502-3p OE vs scramble control; miR-502-3p sponge vs scramble control; miR-502-3p OE vs miR-502-3p sponge) were run for the gene ontology analysis. The top 20 pathways were determined by using the FDR cutoff (0.05) and sorted by fold enrichment. The several biological components-KEGG pathways, cellular process and molecular components were accessed in the analysis.

### Statistical analysis

The statistical analyses of data points were performed using the student’s t-test for analyzing two groups-scramble control vector versus miR-502-3p OE vector, scramble control versus miR-502-3p suppression sponge vector and miR-502-3p OE vector versus miR-502-3p suppression sponge vector. One-way analysis of variance was used for analyzing the results between three groups of samples such as scramble control vector versus miR-502-3p OE vector versus miR-502-3p suppression sponge vector. All the statistical parameters were calculated using GraphPad Prism software, v6 (GraphPad, San Diego, CA, USA) (www.graphpad.com). The P values <0.05 were considered statistically significant.

## RESULTS

### MiR-502-3p modulates the GABRA1 expression

To determine the impact of miR-502-3p on GABRA1, we studied the miR-502-3p and GABRA1 expressions quantitatively by qRT-PCR and qualitatively by RNAScope *in-situ* hybridization analysis. Cells transfected with scramble control vector, miR-502-3p OE vector and miR-502-3p sponge vector displayed robust green fluorescent protein expression as shown in **Figure 1A**. Next, qRT-PCR analysis showed the significantly (P<0.0001) elevated expression of miR-502-3p in OE vector treated cells and suppression (P<0.0001) in sponge vector treated cells relative to control. Opposite to this GABRA1 expression were reduced significantly (P<0.01) in response to OE vector while increase significantly (P<0.01) in sponge treated cells relative to control (**Figure 1B**). Further, miRNA *in-situ* hybridization co-detection analysis were performed using specific probes for hsa-miR-502-3p, U6 snRNA and specific antibody for GABRA1 protein. The GABRA1 stained with LS Duplex green was detected by bright field microscopy. The cells showed increased expression of miR-502-3p probe (red color) and reduced level of GABRA1 protein (blue color) in miR-502-3p OE vector treated cells (**Figure 1A**). U6snRNA used an internal control did not show any change in the expression intensity across experimental conditions. The quantification of miR-502-3p and GABRA1 probe intensity showed that miR-502-3p level was significantly (P<0.0001) increased by miR-502-3p OE vector, opposite to the GABRA1 level decreased (P<0.001) in miR-502-3p OE vector treated cells. Interestingly, GABRA1 level was highly (P<0.0001) increased in miR-502-3p sponge treated cells relative to miR-502-3p OE treated and control (**Figure 1C**). Apparently, miR-502-3p and GABRA1 expression levels were inversely correlated with each other.

**Figure 1.**
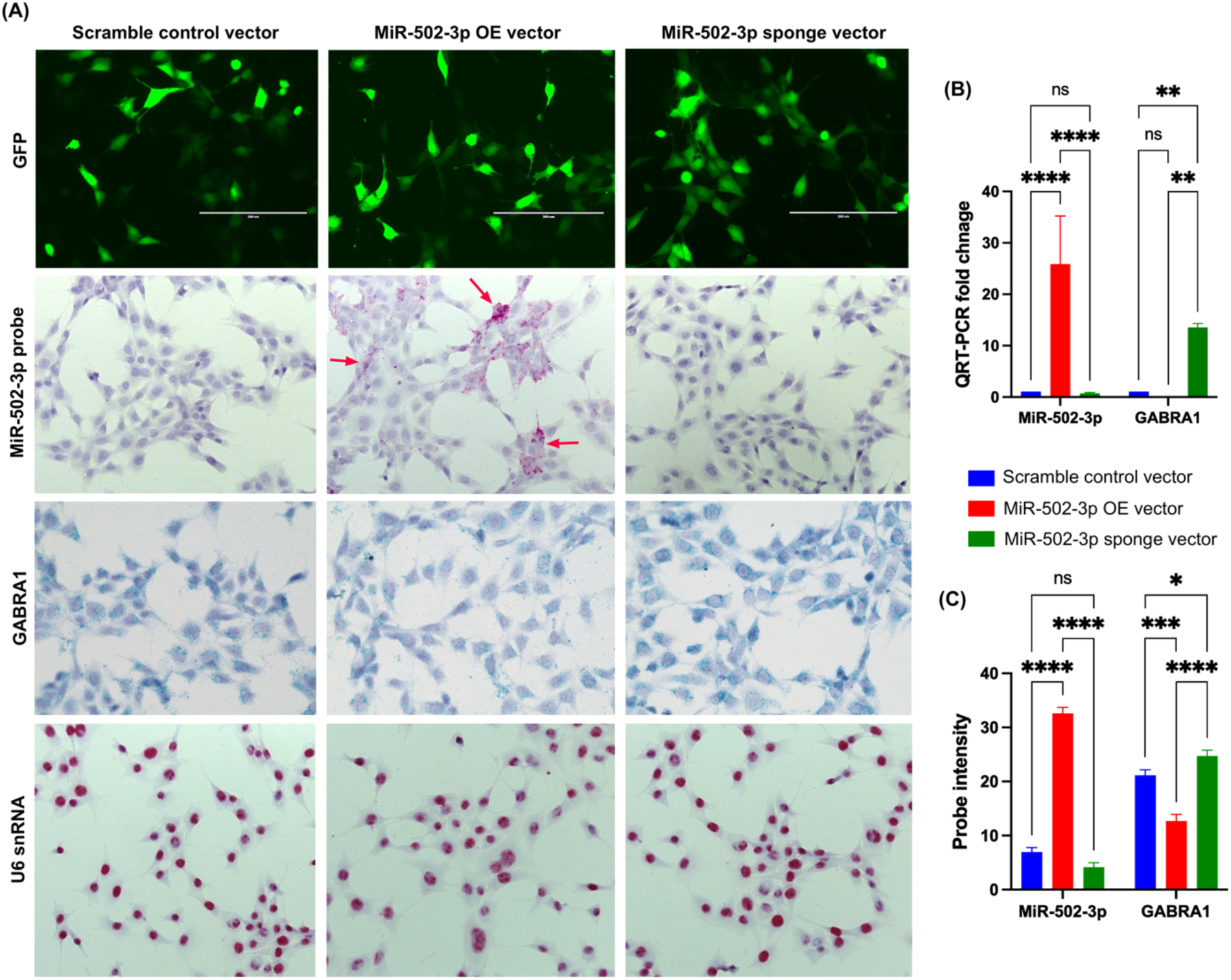
Co-detection of miR-502-3p and GABRA1 expression in HT22 cells. **(A)** Fluorescence microscopy images showing GFP fluorescence in HT22 cells transfected with scramble control vector, miR-502-3p OE vector and miR-502-3p sponge vector. The miRNA *in-situ* hybridization co-detection analysis revealed the localization of miR-502-3p and GABRA1. MiR-502-3p expression was detected using a specific probe (red arrow), and GABRA1 was visualized with LS duplex green staining. The overexpression of miR-502-3p led to a notable decrease in GABRA1 expression compared to the scramble control and sponge-transfected cells. U6 snRNA, used as an internal control, displayed consistent expression across all experimental conditions. **(B)** Quantitative RT-PCR analysis of miR-502-3p expression and GABRA1 mRNA expression in transfected HT22 cells. The miR-502-3p OE plasmid significantly increased miR-502-3p levels (25.90-fold; P <0.0023) compared to the scramble control. MiR-502-3p overexpression resulted in a significant reduction of GABRA1 mRNA levels (∼0.12-fold; P = 0.081), while miR-502-3p sponge transfection led to a significant increase (∼13.56-fold; P < 0.0001) in GABRA1 mRNA expression compared to the scramble control. **(C)** MiR-502-3p probe intensity was significantly increased, whereas GABRA1 intensity was significantly reduced in miR-502-3p OE vector treated cells. Interestingly, GABRA1 intensity was significantly increased in sponge treated cells. Error bars represent standard deviation across all groups. (*P<0.05, **P<0.01, ***P<0.001, ****P< 0.0001).

### Regulation of GABA A receptor and synaptic protein levels by miR-502-3p

To determine the impact of miR-502-3p on GABA A receptor and synaptic proteins, cells were transfected with scramble control vector, miR-502-3p OE vector and miR-502-3p sponge vector and immunoblotting analysis were performed for GABA A subunit proteins (GABRA1, GABRB3, GABRG2, Gephyrin and Calmodulin) and key synaptic proteins (SNAP25, syntaxin1, MAP2, NRXN1 and VAMP2). **Figure 2A** showed the immunoblots for above mentioned proteins in scramble control, miR-502-3p OE and miR-502-3p sponge treated cells. The levels of GABA A receptor and synaptic proteins were found to be reduced by overexpression of miR-502-3p and their levels increased by suppression of miR-502-3p. The densitometry analysis showed the significant upregulation of GABRA1, GABRG2, Gephyrin and Calmodulin proteins in response to miR-502-3p suppression (**Figure 2B**). However, we did not see any significant change in the GABRB3 level. Similarly, the levels of synaptic proteins were also reduced significantly by high level of miR-502-3p and increased by the suppression of miR-502-3p (**Figure 2C**). We did not see any significant change in Syntaxin1 protein level. Together, these data show that high level of miR-502-3p negatively modulate the GABA and synaptic proteins displayed deleterious effects on the cells whereas suppression of miR-502-3p has a positive impact on the GABA and synaptic protein thus could have beneficial role in the neuron function and synaptic activity.

**Figure 2.**
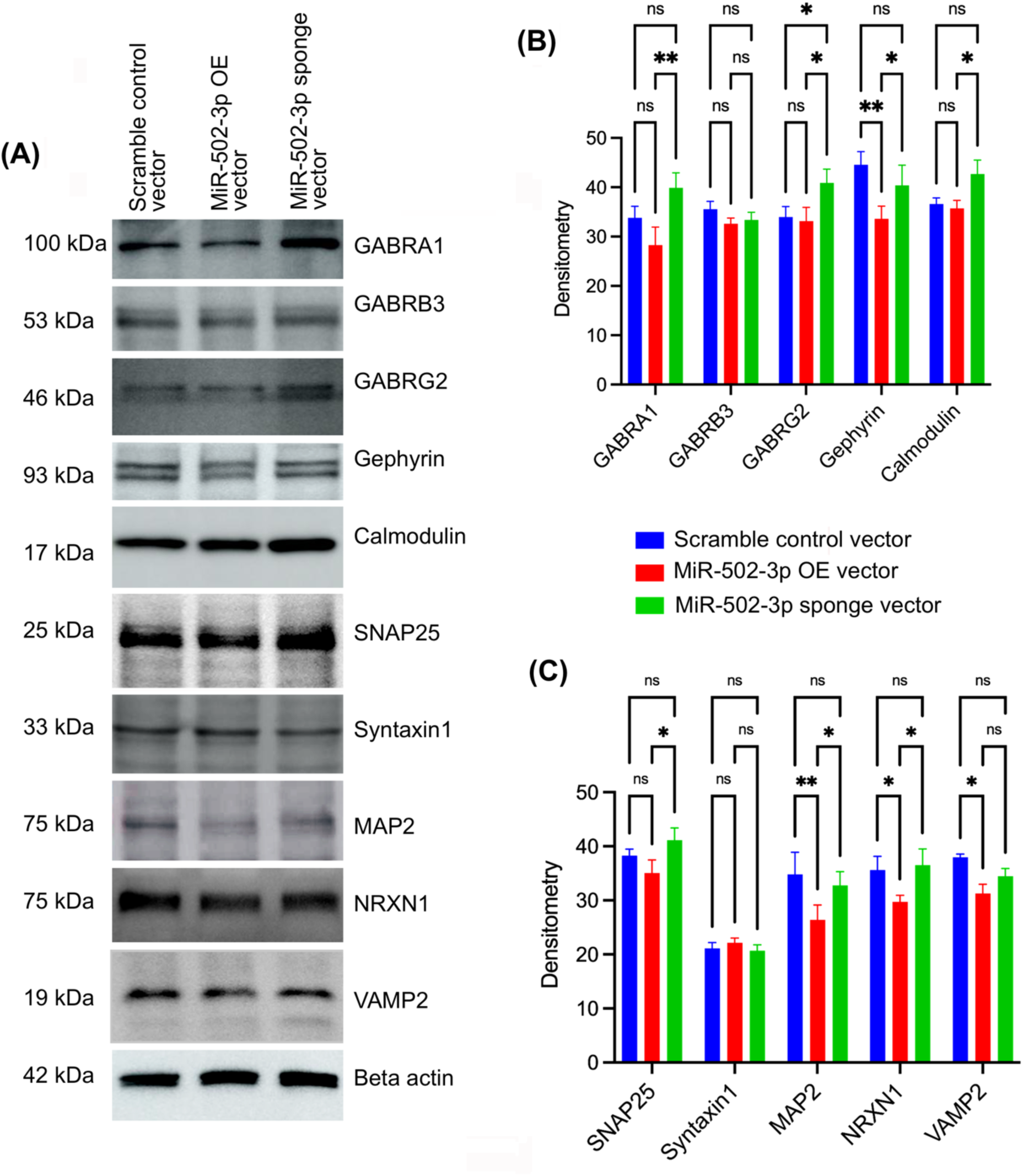
Regulation of GABA A receptor subunits and synaptic protein levels by miR-502-3p in HT22 cells. **(A)** Immunoblotting analysis of GABA A receptor subunit proteins (GABRA1, GABRB3, GABRG2, Gephyrin and Calmodulin) and synaptic proteins (SNAP25, Syntaxin1, MAP2, NRXN1 and VAMP2) in HT22 cells transfected with scramble control vector, miR-502-3p OE vector and miR-502-3p sponge vector. MiR-502-3p overexpression led to a marked reduction in the levels of both GABA A receptor subunits and synaptic proteins compared to the control and sponge-transfected cells. **(B)** Densitometry analysis of immunoblots showing significant reductions in GABRA1, GABRG2, Gephyrin and Calmodulin protein levels in response to miR-502-3p overexpression. **(C)** Densitometry analysis of synaptic proteins, demonstrating significantly reduced levels of MAP2, NRXN1 and VAMP2 proteins in miR-502-3p overexpressed cells relative to the scramble control. Conversely, SNAP25, MAP2 and NRXN1 protein levels were significantly increased in miR-502-3p suppressed cells compared to the scramble control. Error bars represent standard deviation across all groups. (*P<0.05, **P<0.01).

### Effect of miR-502-3p on mitochondrial morphology

MiR-502-3p is a synapse specific miRNA and high content of mitochondria are present at synapse. So, to determine the impact of miR-502-3p on mitochondrial morphology and mitochondrial numbers, TEM analysis was performed on the cells transfected with scramble control vector, miR-502-3p OE vector and miR-502-3p sponge vector. Cells were processed to examine the ultrastructural changes in mitochondrial morphology (**Figure 3A**). TEM images showed that mitochondria appear healthy and elongated in control and sponge treated cells, while mitochondrial quality (size and length) was compromised in miR-502-3p OE treated cells. Mitochondrial number counting showed significant increase in mitochondria number in miR-502-3p OE treated cells relative to control (P<0.01) and sponge treated cells (P<0.001) (**Figure 3B**). Further, mitochondria length (nm) quantification showed the significant reduction in total length of mitochondria in miR-502-3p OE vector treated cells relative to control (P<0.05) and sponge treated cells (P< 0.0001) (**Figure 3C**). These observations confirmed that miR-502-3p affect mitochondria quality, especially ectopic expression of miR-502-3p having an adverse effect on mitochondria morphology while suppression of miR-502-3p improves the mitochondrial quality.

**Figure 3.**
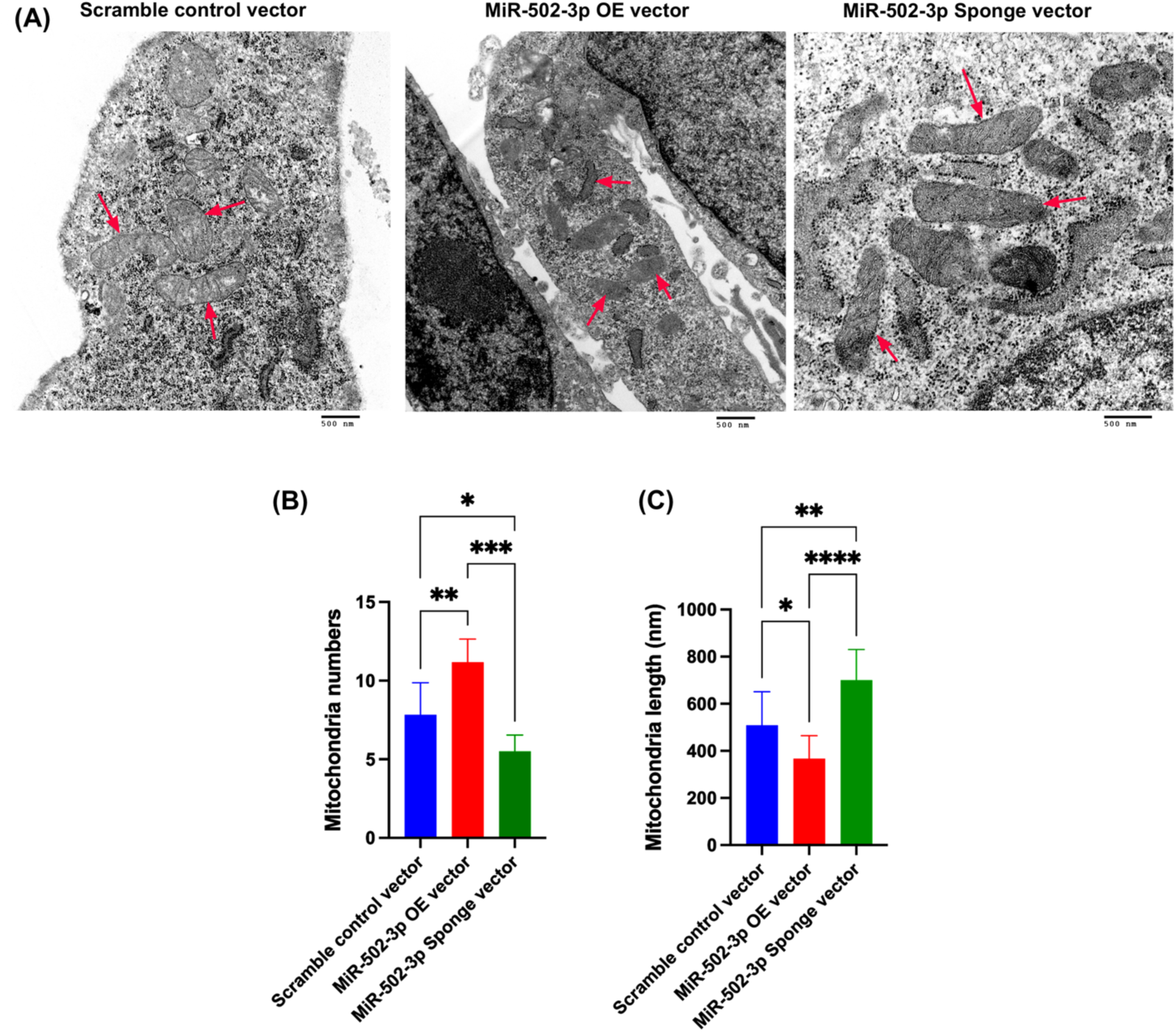
Mitochondria morphology alteration by miR-502-3p. **(A)** Electron microscopy analysis of HT22 cells transfected with scramble control, miR-502-3p OE vector and miR-502-3p sponge vector. The image shows abnormal and fragmented mitochondria in cells transfected with the miR-502-3p OE vector (red arrow), while mitochondria appear healthy and elongated in miR-502-3p sponge transfected cells (red arrow). **(B)** Quantitative analysis of mitochondrial number in the cells transfected with the controls, miR-502-3p OE vector and miR-502-3p sponge treated cells. The graph indicates that number of small and fragmented mitochondria was significantly increased in miR-502-3p OE cells relative to control and sponge treated cells. **(C)** Mitochondria length was significantly reduced in miR-502-3p OE cells while length was significantly increased in sponge treated cells relative to OE and control treated cells. Error bars represent standard deviation across all groups. (*P<0.05, **P<0.01, ***P<0.001, ****P< 0.0001).

### Impact of miR-502-3p on hippocampal neuron genome – Gene array analysis

To determine the genome wide impact of miR-502-3p on hippocampal neurons, cells were transfected with scramble control vector, miR-502-3p OE vector and miR-502-3p sponge vectors in two technical duplicates. Total RNA was isolated from the cells and processed for complete genome analysis using Affymetrix gene array via Clariom S mouse platform. **Table 1** shows a brief summary of number of multiple complex genes, coding genes, non-coding genes and pseudogenes deregulated in three different comparisons. The data were analyzed using three different comparisons-

**Table 1.** Number of coding, non-coding genes deregulated in three different comparisons.

| **Comparison 1- MiR-502-3p OE vector vs Scramble control vector** |  |  |  |  |
| --- | --- | --- | --- | --- |
| **Group** | **Total** | **Passed Filter** | **Up-Regulated** | **Down-Regulated** |
| Multiple_Complex | 11871 | 422 | 191 | 231 |
| Coding | 10261 | 227 | 126 | 101 |
| NonCoding | 60 | 1 | 1 | 0 |
| Pseudogene | 14 | 0 | 0 | 0 |
| All together | 22206 | 650 | 318 | 332 |
| **Comparison 2- MiR-502-3p sponge vector vs Scramble control vector** |  |  |  |  |
| **Group** | **Total** | **Passed Filter** | **Up-Regulated** | **Down-Regulated** |
| Multiple_Complex | 11871 | 476 | 201 | 275 |
| Coding | 10261 | 250 | 132 | 118 |
| NonCoding | 60 | 0 | 0 | 0 |
| Pseudogene | 14 | 0 | 0 | 0 |
| Altogether | 22206 | 726 | 333 | 393 |
| **Comparison 3- MiR-502-3p OE vector vs MiR-502-3p sponge vector** |  |  |  |  |
| **Group** | **Total** | **Passed Filter** | **Up-Regulated** | **Down-Regulated** |
| Coding | 10261 | 62 | 24 | 38 |
| Multiple_Complex | 11871 | 44 | 20 | 24 |
| NonCoding | 60 | 0 | 0 | 0 |
| Pseudogene | 14 | 0 | 0 | 0 |
| Altogether | 22206 | 106 | 44 | 62 |

### (i) MiR-502-3p OE vector versus scramble control vector

To study the change in gene expression profile by miR-502-3p overexpression, gene array analysis was compared between scramble control and miR-502-3p OE treated cells. A total of 650 genes were found to be significantly (P<0.05; fold change >+2 and <-2) deregulated in miR-502-3p overexpressed cells relative to control (**Supplementary Table 1**). Out of them, 318 genes were upregulated and 332 genes we downregulated in miR-502-3p OE cells relative to scramble controls (**Table 1**). Further, we narrowed down the gene selection criteria by fold-change more than +5 and −5 with high significant p-value and FDR p-value. The top 15 most deregulated genes identified with this analysis, with clear differential expression between the miR-502-3p OE and scramble control groups were depicted by heatmap (**Figure 4A**). The fold changes of the top 15 deregulated genes range from −12.65 to +23.27, with corresponding p-values and FDR values indicating statistical significance (**Table 2**). Several genes, including *Spon2*, *Ifit1*, and *Ccl5*, were significantly downregulated, while others like *Prl2c2*, *Prl2c3*, and *Hmox1* were upregulated in response to miR-502-3p OE relative to controls. Notably, deregulated genes were involved in immune response (*Ccl5*, *Ifit1*), oxidative stress (*Hmox1*), and receptor signaling (*Il1rl1*, *Tnfrsf12a*) were significantly altered, reflecting potential pathways influenced by miR-502-3p. A volcano plot illustrates the overall distribution of differentially expressed genes, with a number of transcripts falling below the −log10 (p-value) cutoff of 1.96 (**Figure 4B**). The scatter plot of gene expression profiles showed a clear segregation of deregulated genes between miR-502-3p OE and scramble control conditions (**Figure 4C**).

**Figure 4.**
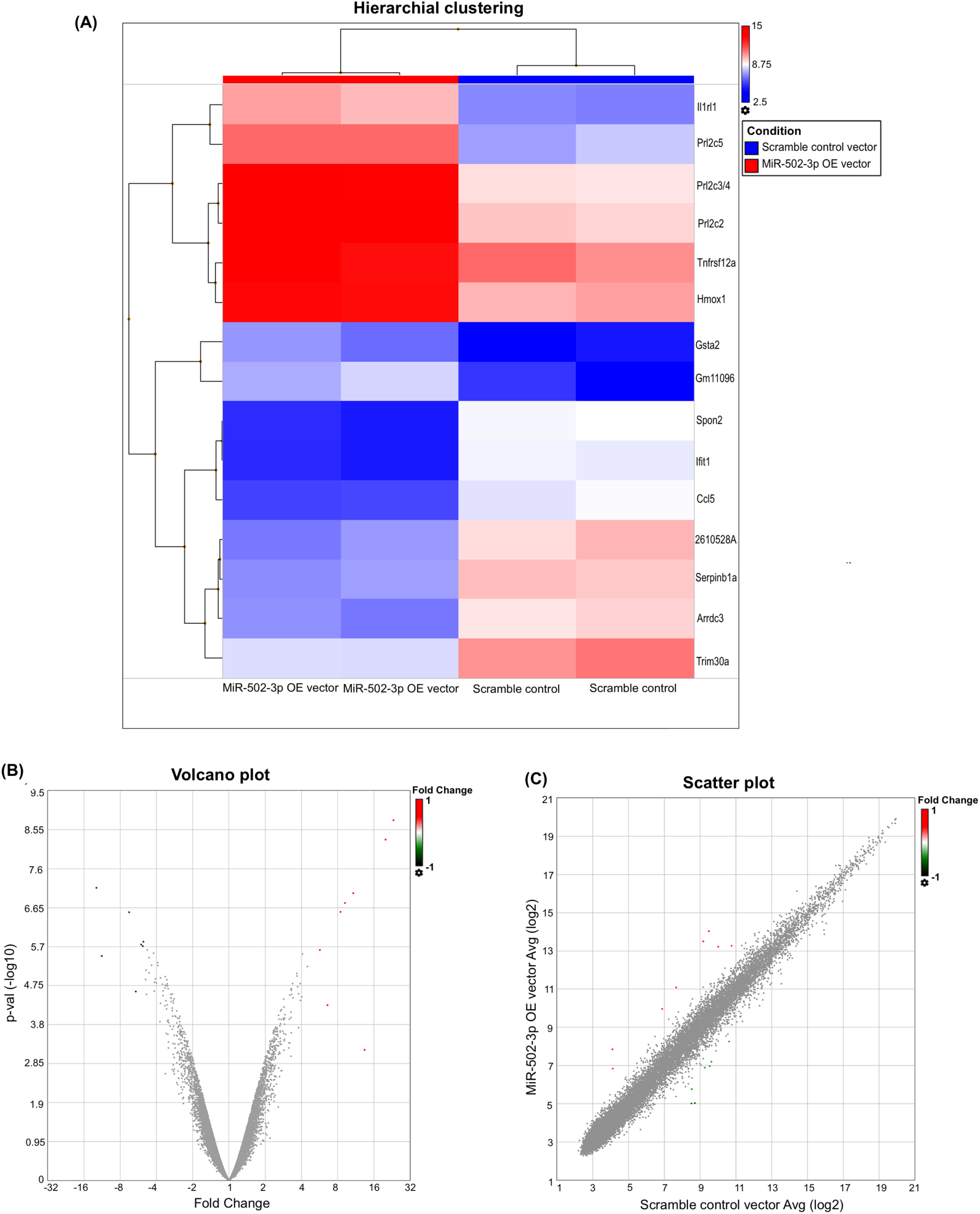
Impact of miR-502-3p overexpression on gene expression profile in HT22 cells. **(A)** Heatmap of the top 15 most deregulated genes identified by microarray analysis comparing HT22 cells transfected with the miR-502-3p OE vector and a scramble control vector. The heatmap illustrates significant differential expression, with fold changes ranging from −12.65 to 23.27. Genes such as Spon2, Ifit1, and Ccl5 are notably down-regulated, whereas Prl2c2, Prl2c3, and Hmox1 are up-regulated. The genes involved in immune response, oxidative stress, and receptor signaling are highlighted to reflect potential pathways influenced by miR-502-3p. Statistical significance is indicated with corresponding p-values and false discovery rate (FDR) values provided in **Table 2. (B)** Volcano plot displaying the overall distribution of differentially expressed genes between miR-502-3p OE and scramble control conditions. The plot shows that several transcripts fall below the −log10 (p-value) cutoff of 1.96, indicating significant differences in gene expression. **(C)** Scatter plot of miRNA expression profiles illustrating the segregation of deregulated genes between miR-502-3p OE and scramble control conditions. The plot demonstrates a clear separation between the two conditions, highlighting the impact of miR-502-3p overexpression on gene expression patterns. (Red color intensity showed upregulated genes and blue color intensity showed downregulated genes).

**Table 2.**
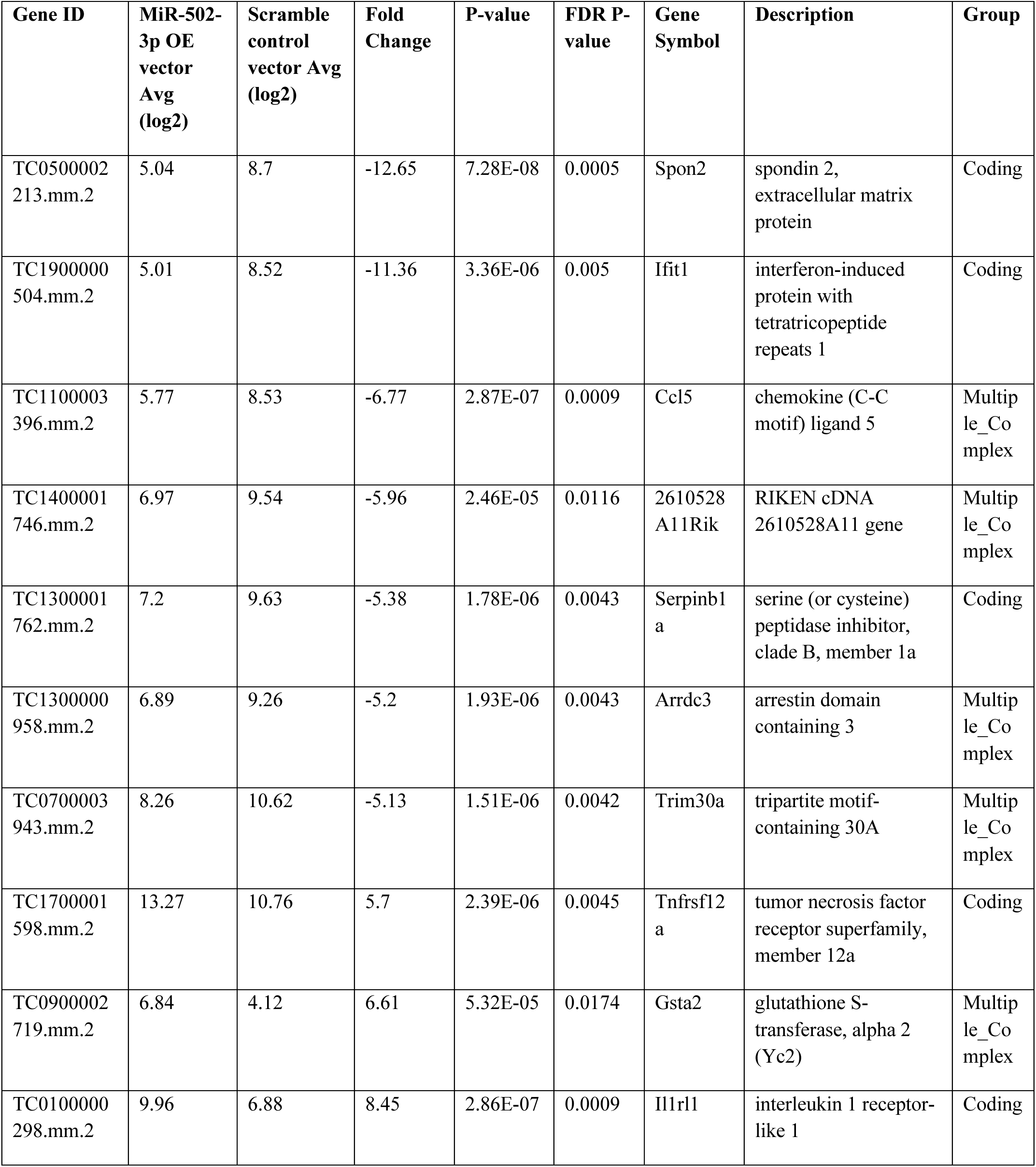
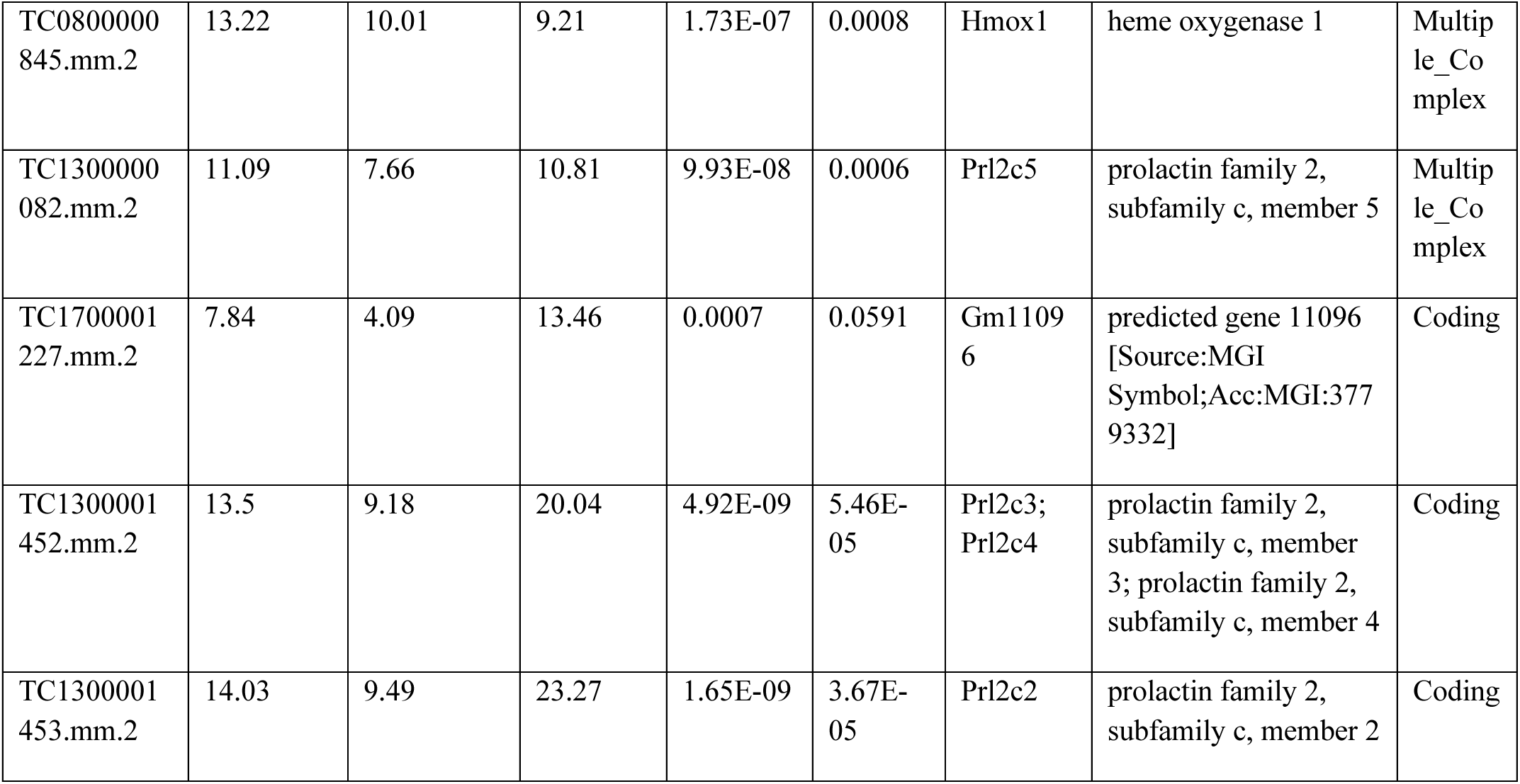
MiR-502-3p OE vector versus Scramble control-most significantly deregulated genes.

### (ii) MiR-502-3p sponge vector versus scramble control vector

To investigate the effect of miR-502-3p inhibition on gene expression profile, we performed the gene array data analysis comparing between the cells transfected with miR-502-3p sponge vector and scramble control vector. A total of 726 genes were found to be significantly deregulated (P<0.05; fold change >+/-2) in miR-502-3p sponge treated cells compared to control (**Supplementary Table 2**). Out of them, 333 genes were significantly upregulated while 393 genes were downregulated by miR-502-3p suppression relative to controls (**Table 1**). Further, we sorted out the top gene by selecting fold-change more than +5 and −5 with high significant p-values and FDR p-values. The heatmap showing the top 13 most deregulated genes identified in the analysis, illustrating marked differences in expression levels between the miR-502-3p sponge and scramble control groups (**Figure 5A**). The fold changes of these top 13 deregulated genes range from −13 to 25.39, with significant p-values and FDR-adjusted p-values (**Table 3**). Several genes such as *Spon2*, *Ifit1*, *Ccl5*, and *Arrdc3* were significantly downregulated upon miR-502-3p inhibition, consistent with their potential role in extracellular matrix remodeling and immune response pathways. Conversely, genes like *Prl2c2*, *Prl2c3*, *Hmox1*, and *Il1rl1* showed substantial upregulation, suggesting their involvement in compensatory mechanisms or alternate signaling pathways triggered by miR-502-3p sponge treatment. The volcano plot depicting the distribution of differentially expressed genes, where several transcripts surpass the −log10(p-value) threshold, indicating strong deregulation (**Figure 5B**). The scatter plot of miRNA expression profiles, with clear divergence between miR-502-3p sponge and scramble control conditions, confirmed the impact of miR-502-3p on the expression of key regulatory genes (**Figure 5C**).

**Figure 5.**
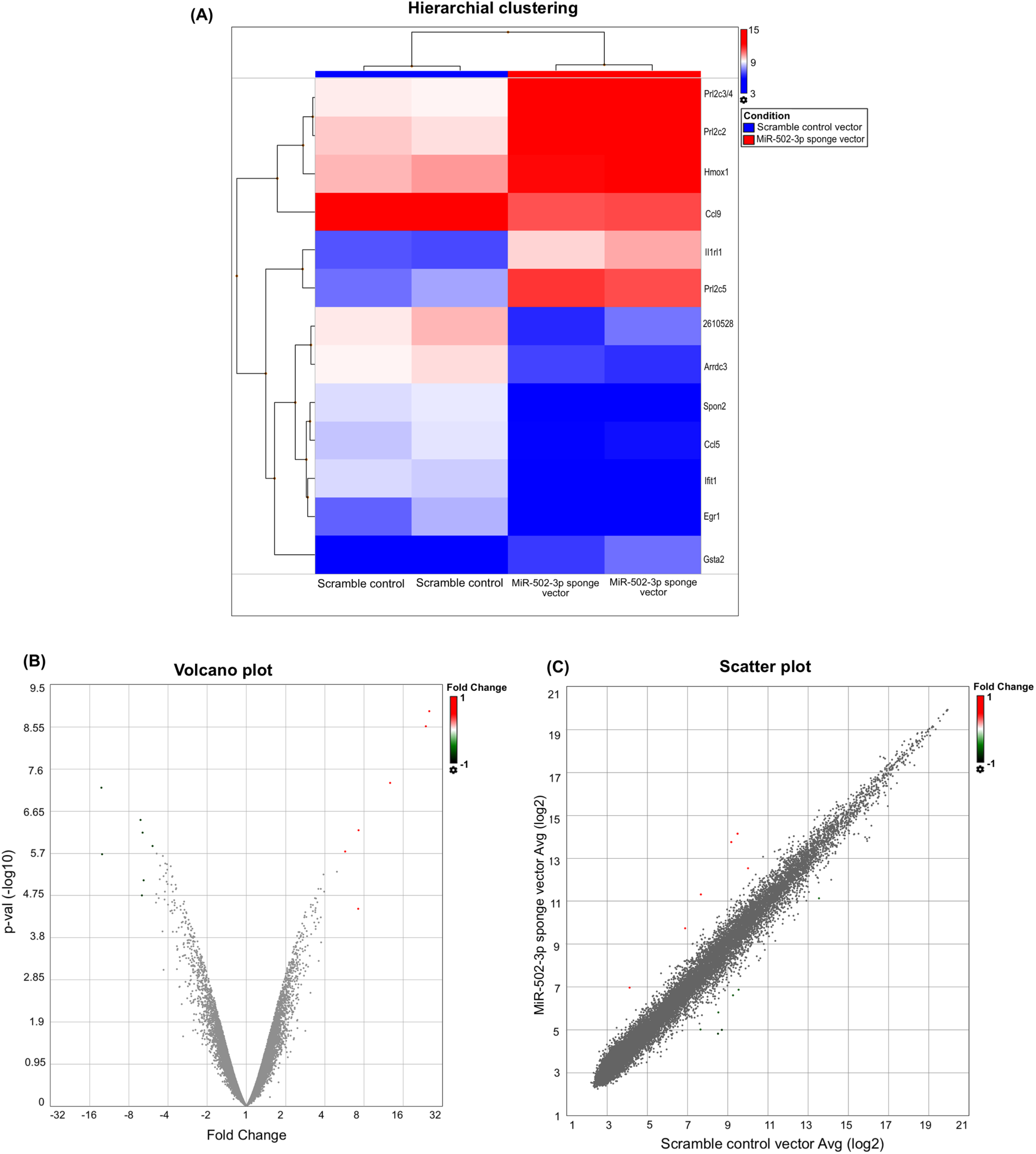
Effect of miR-502-3p inhibition on gene expression profile in HT22 cells. **(A)** Heatmap of the top 13 most deregulated genes identified through transcriptome-wide analysis comparing HT22 cells transfected with the miR-502-3p sponge vector and a scramble control vector. The heatmap highlights marked differences in gene expression levels, with fold changes ranging from −13 to 25.39. Key genes such as Spon2, Ifit1, Ccl5, and Arrdc3 are significantly down-regulated, whereas Prl2c2, Prl2c3, Hmox1, and Il1rl1 are notably up-regulated. These expression changes reflect potential roles in extracellular matrix remodeling, immune response pathways, and compensatory mechanisms. Statistical significance is provided with p-values and FDR-adjusted p-values detailed in **Table 3**. **(B)** Volcano plot showing the distribution of differentially expressed genes between miR-502-3p sponge and scramble control conditions. The plot illustrates that several transcripts surpass the −log10(p-value) threshold, indicating significant deregulation and highlighting the impact of miR-502-3p inhibition on gene expression. **(C)** Scatter plot of miRNA expression profiles demonstrating clear separation between miR-502-3p sponge and scramble control conditions. This segregation confirms the significant effect of miR-502-3p inhibition on the expression of key regulatory genes. (Red color intensity showed upregulated genes and blue color intensity showed downregulated genes).

**Table 3.** MiR-502-3p sponge versus Scramble control - most significantly deregulated genes.

| Gene ID | MiR-502-3p sponge vector Avg (log2) | Scramble control vector Avg (log2) | Fold Change | P-value | FDR P-value | Gene Symbol | Description | Group |
| --- | --- | --- | --- | --- | --- | --- | --- | --- |
| TC0500002213.mm.2 | 5 | 8.7 | -13 | 6.54E-08 | 0.0004 | Spon2 | spondin 2, extracellular matrix protein | Coding |
| TC1900000504.mm.2 | 4.83 | 8.52 | -12.9 | 2.07E-06 | 0.0042 | Ifit1 | interferon-induced protein with tetratricopeptide repeats 1 | Coding |
| TC1100003396.mm.2 | 5.82 | 8.53 | -6.52 | 3.49E-07 | 0.0015 | Ccl5 | chemokine (C-C motif) ligand 5 | Multiple_Co_mplex |
| TC1400001746.mm.2 | 6.87 | 9.54 | -6.37 | 1.77E-05 | 0.0104 | 2610528A11Rik | RIKEN cDNA 2610528A11 gene | Multiple_Co_mplex |
| TC1300000958.mm.2 | 6.62 | 9.26 | -6.27 | 6.83E-07 | 0.0022 | Arrdc3 | arrestin domain containing 3 | Multiple_Co_mplex |
| TC1800000301.mm.2 | 5.02 | 7.65 | -6.16 | 7.98E-06 | 0.0071 | Egr1 | early growth response 1 | Coding |
| TC1100003397.mm.2 | 11.14 | 13.54 | -5.26 | 1.36E-06 | 0.0038 | Ccl9 | chemokine (C-C motif) ligand 9 | Multiple_Co_mplex |
| TC0800000845.mm.2 | 12.53 | 10.01 | 5.72 | 1.82E-06 | 0.0042 | Hmox1 | heme oxygenase 1 | Multiple_Co_mplex |
| TC0900002719.mm.2 | 6.97 | 4.12 | 7.24 | 3.53E-05 | 0.0135 | Gsta2 | glutathione S-transferase, alpha 2 (Yc2) | Multiple_Co_mplex |
| TC0100000298.mm.2 | 9.74 | 6.88 | 7.25 | 5.96E-07 | 0.0022 | Il1rl1 | interleukin 1 receptor-like 1 | Coding |
| TC130000008<br>2.mm.2 | 11.32 | 7.66 | 12.69 | 5.16E-08 | 0.0004 | Prl2c5 | prolactin family<br>2, subfamily c,<br>member 5 | Multip<br>le_Co<br>mplex |
| TC130000145<br>2.mm.2 | 13.76 | 9.18 | 23.87 | 2.74E-09 | 3.05E-05 | Prl2c3;<br>Prl2c4 | prolactin family<br>2, subfamily c,<br>member 3;<br>prolactin family<br>2, subfamily c,<br>member 4 | Codin<br>g |
| TC130000145<br>3.mm.2 | 14.15 | 9.49 | 25.39 | 1.24E-09 | 2.76E-05 | Prl2c2 | prolactin family<br>2, subfamily c,<br>member 2 | Codin<br>g |

### (iii) MiR-502-3p OE vector versus miR-502-3p sponge vector

Lastly, to investigate the gene expression changes in miR-502-3p overexpression versus inhibition, we performed a comparative transcriptomic analysis between cells transfected with a miR-502-3p OE vector and those with a miR-502-3p sponge vector. A total of 106 genes were found to be significantly (P<0.05; fold change >+/-2) deregulated in this comparison as mentioned in **Supplementary Table 3**. The 44 genes were upregulated, and 72 genes were significantly downregulated by miR-502-3p overexpression relative to miR-502-3p suppression (**Table 1**). Next, we screened top deregulated genes by selecting the fold-change more than +2.4 and −2.4 with high significant p-values and FDR p-values. The heatmap in **Figure 6A**/ shows the top 20 most deregulated genes identified in this comparison, highlighting distinct expression patterns associated with miR-502-3p modulation. The fold changes of these top 20 deregulated genes, revealing fold changes ranging from −2.93 to 16.93 (**Table 4**). While the unadjusted p-values suggest statistical significance for many of these genes (all P-value < 0.05), the false discovery rate (FDR P-value) for all entries is 0.8678. This high FDR indicates that, after adjusting for multiple comparisons, these changes may not be statistically significant. Nonetheless, the substantial fold changes observed warrant further exploration to understand their biological implications. Among the downregulated genes in the miR-502-3p OE vector compared to the sponge vector, notable entries include: Spon2 (*spondin 2*), an extracellular matrix protein involved in cell adhesion and migration, with a fold change of −2.93. Atp4a (*ATPase, H⁺/K⁺ exchanging, gastric, alpha polypeptide*), essential for maintaining gastric acidity, with a fold change of −2.69. Ifit1 (*interferon-induced protein with tetratricopeptide repeats 1*), implicated in antiviral responses, with a fold change of −2.64. Conversely, several genes were markedly upregulated in the miR-502-3p OE condition: Prl2c2 (*prolactin family 2, subfamily c, member 2*), with a fold change of 25.39. Prl2c3; Prl2c4 (*prolactin family 2, subfamily c, member 3 and 4*), with a combined fold change of 23.87. Prl2c5 (*prolactin family 2, subfamily c, member 5*), exhibiting a fold change of 12.69. These prolactin family members are involved in various hormonal signaling pathways, suggesting that miR-502-3p may influence endocrine functions within neurons. The distribution of differentially expressed genes, by volcano plot highlighting those with the most significant changes (**Figure 6B**). Scatter plot of gene expression profiles, demonstrating clear separation between the miR-502-3p OE and sponge vector conditions, which corroborates the heatmap findings despite the high FDR values (**Figure 6C**).

**Figure 6.**
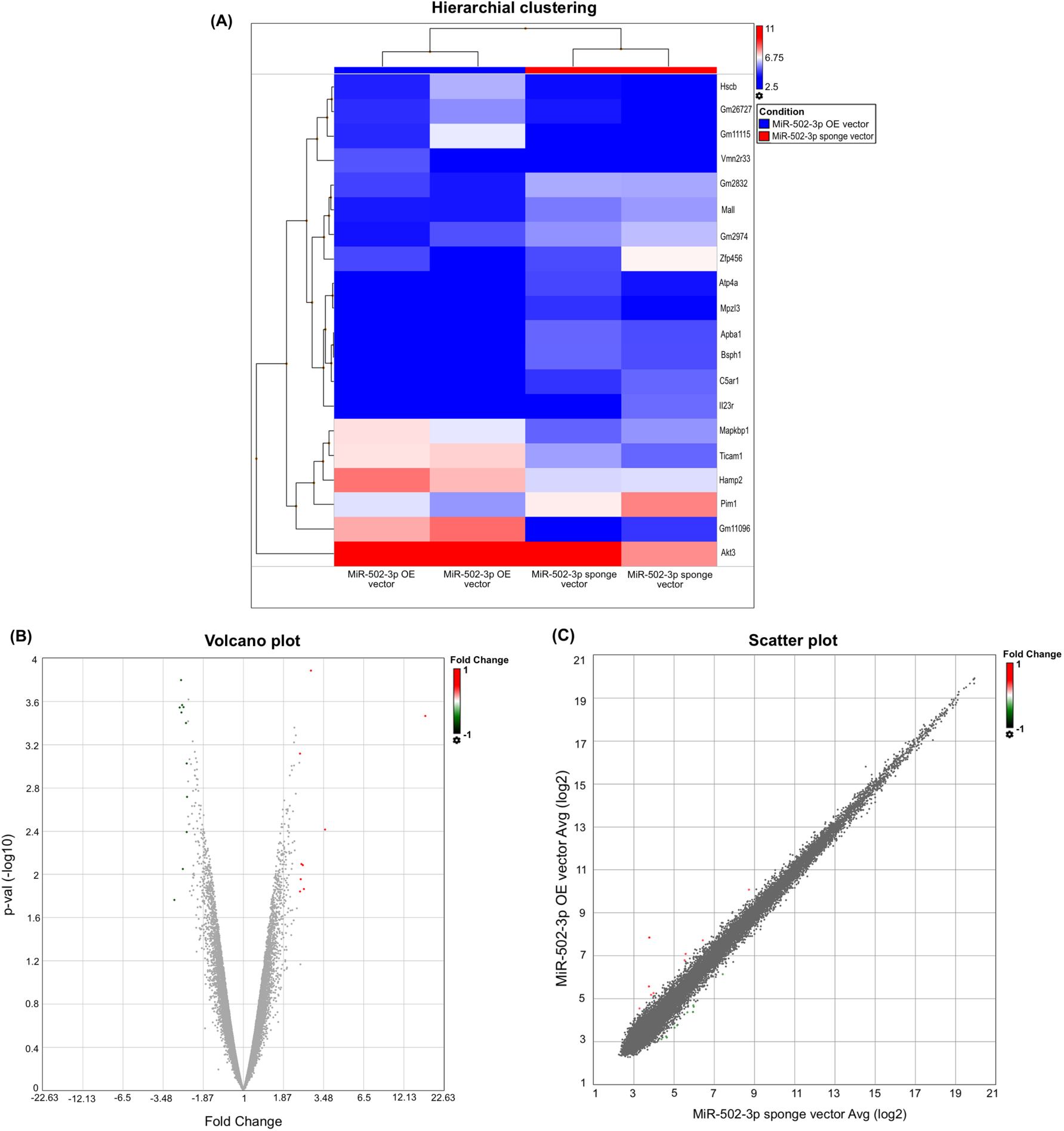
Comparative transcriptomic analysis of miR-502-3p overexpression and inhibition in HT22 Cells. **(A)** Heatmap of the top 20 most deregulated genes from transcriptomic analysis comparing HT22 cells transfected with a miR-502-3p overexpression (OE) vector and a miR-502-3p sponge vector. The heatmap shows distinct expression patterns, with fold changes ranging from −2.93 to 25.39. Notable genes downregulated in the miR-502-3p OE condition include Spon2 (−2.93), Atp4a (−2.69), and Ifit1 (−2.64). Upregulated genes include Prl2c2 (25.39), Prl2c3/Prl2c4 (23.87 combined), and Prl2c5 (12.69). Despite many unadjusted p-values indicating statistical significance (P-val < 0.05), the high false discovery rate (FDR P-val = 0.8678) suggests that these changes may not be statistically significant after multiple comparison adjustments. The data is detailed in **Table 4**. **(B)** Volcano plot illustrating the distribution of differentially expressed genes between miR-502-3p OE and sponge vector conditions. The plot highlights genes with the most significant changes in expression, with several transcripts surpassing the −log10(p-value) threshold, although the high FDR values caution against overinterpretation. **(C)** Scatter plot of gene expression profiles showing clear separation between the miR-502-3p OE and sponge vector conditions. This segregation confirms the differential expression patterns observed in the heatmap, underscoring the substantial impact of miR-502-3p modulation despite high FDR values. (Red color intensity showed upregulated genes and blue color intensity showed downregulated genes).

**Table 4.**
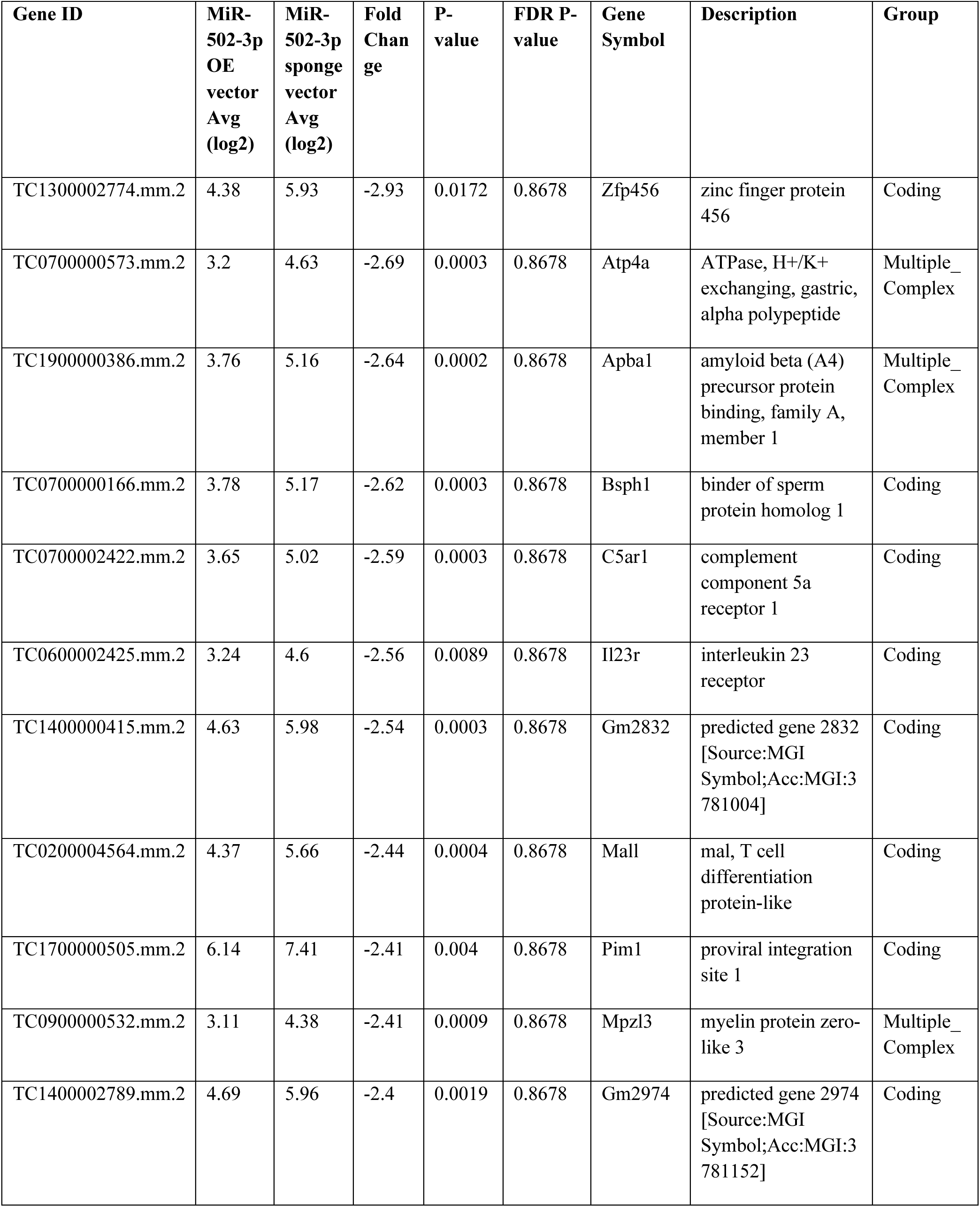
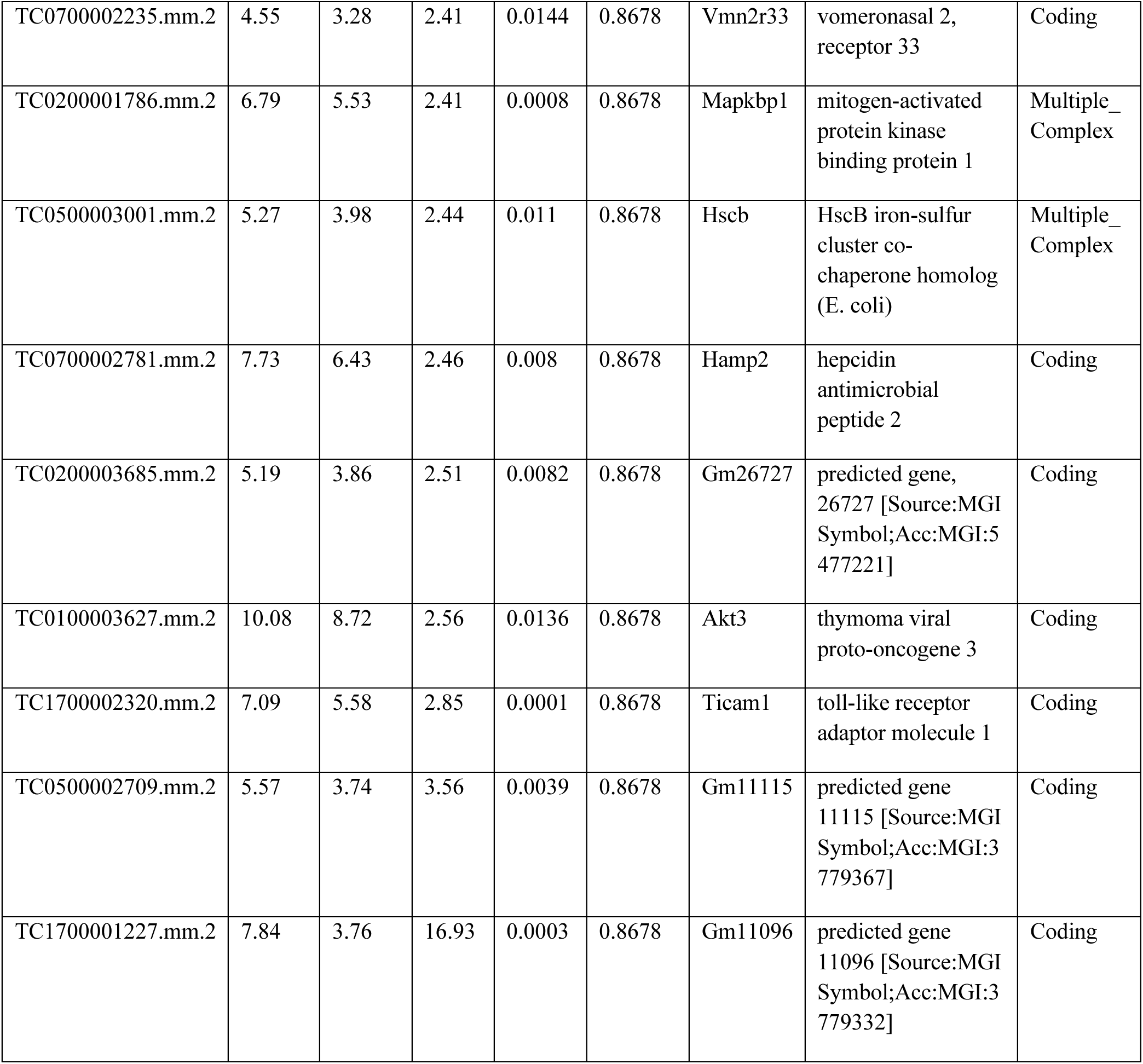
MiR-502-3p OE versus miR-502-3p Sponge - most significantly deregulated genes.

In summary, the modulation of miR-502-3p in hippocampal neurons leads to substantial changes in the expression of genes related to extracellular matrix composition, immune response, and hormonal signaling. While the high FDR values suggest that these findings should be interpreted with caution, the pronounced fold changes observed in key genes indicate that miR-502-3p may play a significant role in regulating critical biological pathways. Future studies with larger sample sizes or alternative validation methods are recommended to confirm these preliminary findings.

### Bioinformatic analysis of miR-502-3p

To understand the biological significance of miR-502-3p in cellular and molecular function of cells, we conducted in-silico gene ontology analysis to deregulated genes. The relevance of ectopic expression of miR-502-3p and deregulated genes were studied in KEGG pathways, cellular process and molecular components. The top cellular pathways affected by miR-502-3p overexpression are relevant to cancer and metabolism (**Figure 7A**). However, brain related pathways such as FoxO signaling, and cellular senescence were also detected in the analysis.

**Figure 7.**
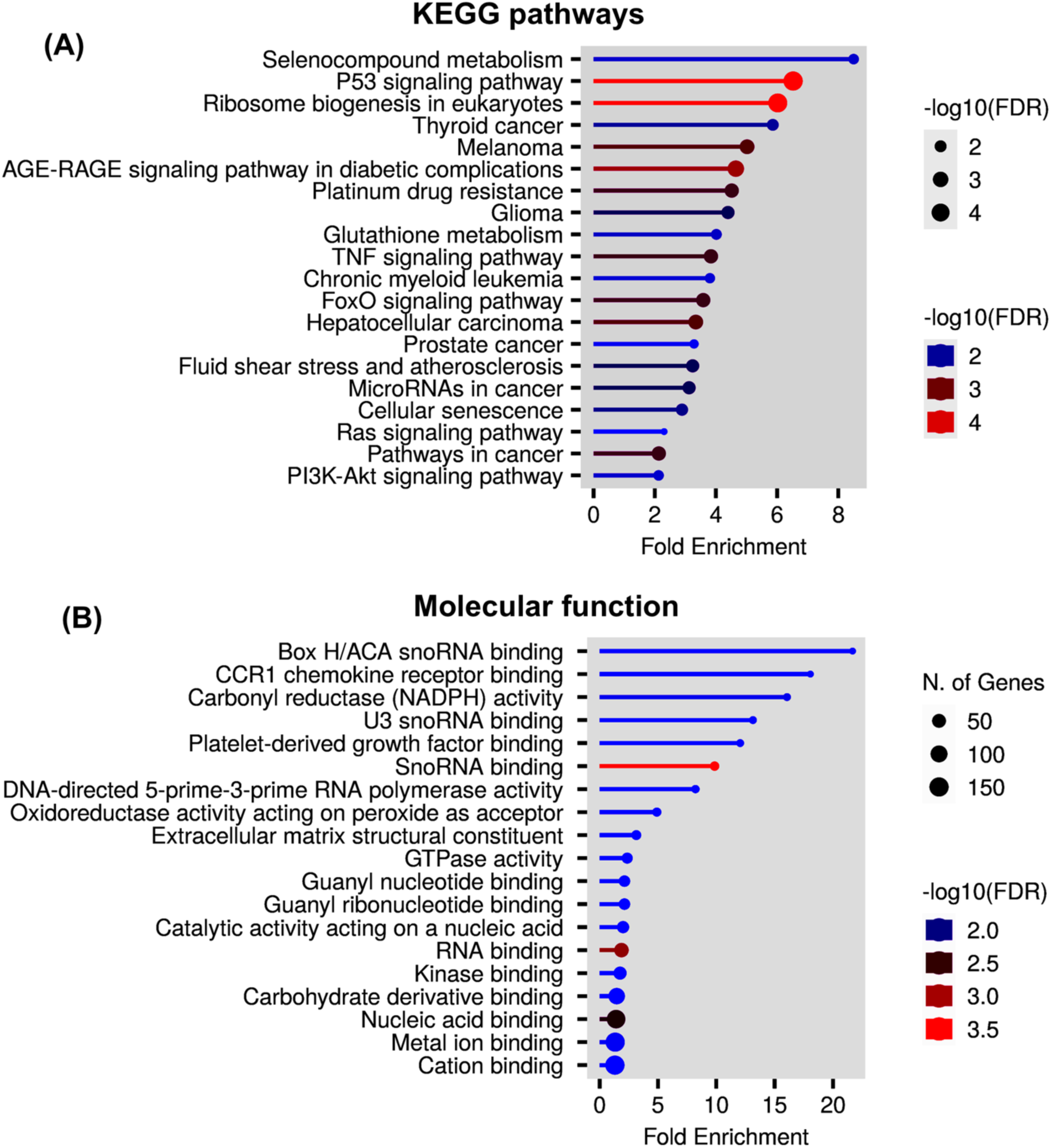
Gene ontology enrichment analysis of miR-502-3p. **(A)** KEGG pathway analysis of deregulated genes in response to miR-502-3p overexpression. **(B)** The topmost molecular function of genes that are deregulated in response to overexpression of miR-502-3p.

Further, molecular function analysis unveiled the significant involvement of miRNAs in several mitochondrial and oxidative stress related pathways (**Figure 7B**). All together in-silico analysis unveiled that overexpression of miR-502-3p significantly impact the cancer related pathways, brain function and mitochondrial activities.

## DISCUSSION

MiRNAs relevance in human disease received the noble recognition in 2024. Several miRNAs have been identified in the human body which are critically associated with cells and tissue specific biological functions (22,23). Each miRNA has special biological characteristics and are critically associated with specific disease(s) (3,24). MiR-502-3p is involved in many human diseases (18), however biological relevance of miR-502-3p has been recently investigated in neurodegenerative disease such as AD (5,16–19). Higher content in AD synapse and synapse localized nature launched miR-502-3p as an interesting molecule worth investigating in AD with focus on synaptic function. As an extension of ongoing lab project, current study was focused to determine the impact of miR-502-3p on GABA subunit proteins, local synaptic proteins, mitochondrial morphology and genome wide impact of miR-502-3p on hippocampal neuron function.

We have studied the role of miR-502-3p on GABAergic synapse function (19), however to examine the miR-502-3p co-localization with GABRA1 protein we re-investigated the impact of miR-502-3p on GABRA1 levels. The qRT-PCR and RNAScope *in-situ* hybridization analysis confirmed an inverse relationship of miR-502-3p with GABRA1 protein. Given that GABRA1 encodes the α1 subunit of the GABA A receptor, which is crucial for mediating inhibitory neurotransmission in the central nervous system (CNS), the suppression of GABRA1 by miR-502-3p may have significant implications for neuronal function. The impact of miR-502-3p on other GABA receptor subunits extended beyond GABRA1, as evidenced by the significant reduction in GABRB3, GABRG2, Gephyrin and Calmodulin protein levels in response to miR-502-3p overexpression. These subunits play essential roles in forming functional GABA A receptors, and their coordinated regulation suggests that miR-502-3p may have a broader influence on GABAergic neurotransmission. Altered GABAergic signaling is associated with various neuropsychiatric disorders, including AD, epilepsy, anxiety, and schizophrenia (5). Thus, the regulatory relationship between miR-502-3p and GABRA1 may provide a potential therapeutic target for conditions characterized by GABAergic dysfunction.

Next, we determined the beneficial and/or deleterious effects of high synapse level of miR-502-3p in relation to synaptic function. The high level of miR-502-3p negatively modulates the key synaptic proteins (SNAP25, Syntaxin1, VAMP2, PSD95). All these proteins play crucial roles in normal synaptic function such as synaptic vesicle docking and neurotransmitter release (25,26) and gets deregulated in various neurological disorders (27,28). Suppressed expression of miR-502-3p augment the levels of these proteins indicating a potential compensatory mechanism that may enhance synaptic function when miR-502-3p activity is diminished. These results highlight the complex regulatory network orchestrated by miR-502-3p, suggesting that its effects are not limited to a single receptor or synaptic protein but encompass broader synaptic processes.

Our study also demonstrated that miR-502-3p overexpression induces significant alterations in mitochondrial morphology, as observed through TEM analysis. Mitochondrial accumulation at the synapse plays critical roles in synaptic energy production, calcium homeostasis, synaptic vesicles turnover-release and recycling and the regulation of apoptosis (29,30), all of which are essential for neuronal function. The mitochondria in miR-502-3p overexpressed cells were smaller and exhibited a distorted structure compared to the elongated mitochondria in miR-502-3p suppressed cells. The shorter mitochondrial length observed in miR-502-3p overexpressed cells is particularly noteworthy, as it may indicate impaired mitochondrial dynamics, including fission and fusion processes. Dysregulation of mitochondrial morphology and function has been implicated in various neurological conditions, including AD and Parkinson’s disease, where disrupted mitochondrial dynamics lead to energy deficits and increased neuronal vulnerability to stress (31–33). The observed mitochondrial abnormalities in miR-502-3p overexpressing cells suggest that miR-502-3p may contribute to mitochondrial dysfunction whereas knockdown of miR-502-3p could have protective effect on mitochondrial function.

One of the major purposes of this study was to examine the other cellular genes that are deregulated in response to elevation and reduction of miR-502-3p. Gene array analysis revealed significant gene expression changes in response to both miR-502-3p overexpression and inhibition. Notably, miR-502-3p overexpression led to the downregulation of genes involved in immune response (e.g., Ifit1, Ccl5), (34–36) while genes related to oxidative stress and endocrine signaling (e.g., Hmox1, Prl2c family) were upregulated (37). These findings suggest that miR-502-3p may influence diverse cellular pathways, beyond its role in regulating GABA receptor expression. Similarly, miR-502-3p inhibition resulted in marked changes in the expression of extracellular matrix components (e.g., Spon2) which is critically associated with amyloid beta production and cognitive impairment in AD (38,39). Further, miR-502-3p over/under expression modulates other significant genes (Atp4a, Apba1 and Mapkbp1) that are associated with AD progression (41,42). Moreover, in-silico bioinformatic analysis unveiled some top KEGG pathways and molecular function of miR-502-3p associated with cellular senescence, oxidative stress and immune response genes, highlighting the potential role of miR-502-3p in modulating cell adhesion, migration, and inflammatory processes. These transcriptomic changes provide a comprehensive overview of the molecular impact of miR-502-3p on neuronal cells and underscore its multifaceted role in cellular homeostasis.

The comprehensive effects of miR-502-3p on GABA A receptor subunits, synaptic proteins, mitochondrial structure, and gene expression underscore its significance in neuronal activity and other cellular function. The suppression of GABA A receptor subunits and synaptic proteins by miR-502-3p suggests that it may play a role in modulating synaptic inhibition and excitability, with potential implications for neurological disorders characterized by altered GABAergic transmission. Furthermore, the mitochondrial abnormalities observed in miR-502-3p overexpressed cells raise the possibility that miR-502-3p may contribute to oxidative stress, synaptic energy deprivation and neuron death in neurodegenerative processes. Future studies should explore the functional consequences of these mitochondrial changes, including their impact on energy metabolism and neuronal survival. The ongoing research projects in our lab are investigating the impact of miR-502-3p on GABA function, synaptic activity, mitochondrial function and cognitive function in AD using the stereotaxic injections of miR-502-3p OE and sponge lentivirus in APP (5XFAD) and Tau (P301L) mouse models.

In conclusion, our findings established miR-502-3p as a key regulator of GABA A receptor subunits, synaptic proteins, and mitochondrial morphology in hippocampal neuron cells. These results provide a foundation for future investigations into the therapeutic potential of miR-502-3p in neurological disorders. More research is warranted to investigate the synapse wide (pre and post synaptic) impact of miR-502-3p in different brain regions and on the different kind of neurons/synapses across AD brain.

## ACKNOWLEDGMENTS

The authors are exceedingly grateful to Prof. Rajkumar Lakshmanaswamy, Chair of the Department of Molecular and Translational Medicine, TTUHSC El Paso for the immense research support. We would like to thank our lab members Ms. Melissa Torres, Ms. Aditi Kulkarni, Mr. Morgan Ogwo, Mr. Davin Devara, Ms. Yogyana and Ms. Angelica.

## AUTHORS CONTRIBUTION

Conceptualization and supervision: SK; experimental performance: BS, SK, DR, GG and SR; analysis, interpretation, and validation of data: SK, BS and DR; writing and original draft preparation: SK and BS; review, editing, and finalization of manuscript: BS, DR, GG, SR and SK. All authors have read and agreed to the published version of the manuscript.

## DATA AVAILABILITY

Data will be made available on request.

