## Supplementary material for "MicroRNA-502-3p modulates the GABA A subunits, synaptic proteins and mitochondrial morphology in hippocampal neurons": SI File 1

### Vector Summary

| | |
|---|---|
| **Vector ID** | VB900088-2245cyx |
| **Vector Name** | pRP[Exp]-CAG>EGFP |
| **Vector Size** | 4801 bp |
| **Vector Type** | Mammalian Gene Expression Vector |
| **Inserted Promoter** | CAG |
| **Inserted ORF** | EGFP |
| **Plasmid Copy Number** | High |
| **Antibiotic Resistance** | Ampicillin |
| **Cloning Host** | Stbl3 (or alternative strain) |

### Vector Map

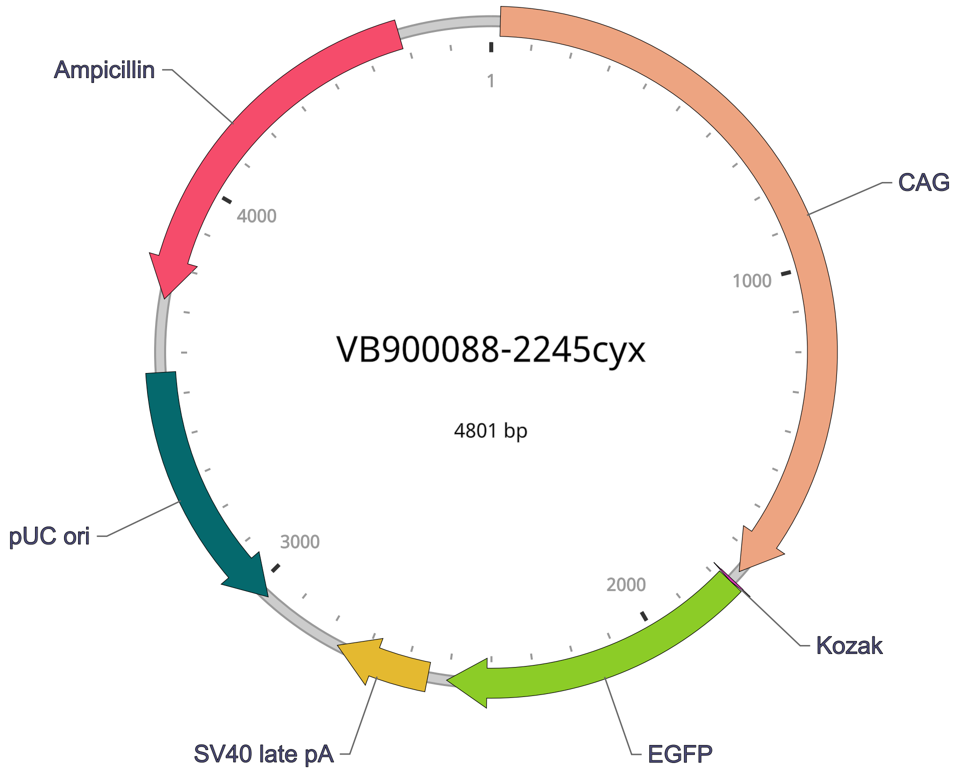


### Vector Components

| Name | Position | Size (bp) | Type | Description | Application notes |
|---|---|---|---|---|---|

| Name | Position | Size (bp) | Type | Description | Application notes |
|---|---|---|---|---|---|
| **CAG** | 22-1754 | 1733 | Promoter | CMV early enhancer fused to modified chicken β-actin promoter | Strong promoter. |
| Kozak | 1779-1784 | 6 | Miscellaneous | Kozak translation initiation sequence | Facilitates translation initiation of ATG start codon downstream of the Kozak sequence. |
| **EGFP** | 1785-2504 | 720 | CDS | Enhanced green fluorescent protein; codon optimized based on a variant of wild type GFP from the jellyfish Aequorea victoria | Commonly used green fluorescent protein; ranked high in brightness, photostability and pH stability among all fluorescent proteins. |
| SV40 late pA | 2549-2770 | 222 | PolyA_signal | Simian virus 40 late polyadenylation signal | Allows transcription termination and polyadenylation of mRNA transcribed by Pol II RNA polymerase. |
| pUC ori | complement (2966-3554) | 589 | Rep_origin | pUC origin of replication | Facilitates plasmid replication in E. coli; regulates high-copy plasmid number (500-700). |
| Ampicillin | complement (3725-4585) | 861 | CDS | Ampicillin resistance gene | Allows E. coli to be resistant to ampicillin. |

**Note:** Components added by user are listed in **bold red** text.

### Vector Sequence

```
  1 CAACTTTGTA TAGAAAAGTT GCTCGACATT GATTATTGAC TAGTTATTAA TAGTAATCAA TTACGGGGTC ATTAGTTCAT
 81 AGCCCATATA TGGAGTTCCG CGTTACATAA CTTACGGTAA ATGGCCCGCC TGGCTGACCG CCCAACGACC CCCGCCCATT
161 GACGTCAATA ATGACGTATG TTCCCATAGT AACGCCAATA GGGACTTTCC ATTGACGTCA ATGGGTGGAG TATTTACGGT
241 AAACTGCCCA CTTGGCAGTA CATCAAGTGT ATCATATGCC AAGTACGCCC CCTATTGACG TCAATGACGG TAAATGGCCC
321 GCCTGGCATT ATGCCCAGTA CATGACCTTA TGGGACTTTC CTACTTGGCA GTACATCTAC GTATTAGTCA TCGCTATTAC
401 CATGGTCGAG GTGAGCCCCA CGTTCTGCTT CACTCTCCCC ATCTCCCCCC CCTCCCCACC CCCAATTTTG TATTTATTTA
481 TTTTTTAATT ATTTTGTGCA GCGATGGGGG CGGGGGGGGG GGGGGGGCGC GCGCCAGGCG GGGCGGGGCG GGGCGAGGGG
561 CGGGGCGGGG CGAGGCGGAG AGGTGCGGCG GCAGCCAATC AGAGCGGCGC GCTCCGAAAG TTTCCTTTTA TGGCGAGGCG
641 GCGGCGGCGG CGGCCCTATA AAAAGCGAAG CGCGCGGCGG GCGGGAGTCG CTGCGCGCTG CCTTCGCCCC GTGCCCCGCT
```

```
721  CCGCCGCCGC CTCGCGCCGC CCGCCCCGGC TCTGACTGAC CGCGTTACTC CCACAGGTGA GCGGGCGGGA CGGCCCTTCT
801  CCTCCGGGCT GTAATTAGCG CTTGGTTTAA TGACGGCTTG TTTCTTTTCT GTGGCTGCGT GAAAGCCTTG AGGGGCTCCG
881  GGAGGGCCCT TTGTGCGGGG GGAGCGGCTC GGGGGGTGCG TGCGTGTGTG TGTGCGTGGG GAGCGCCGCG TGCGGCTCCG
961  CGCTGCCCGG CGGCTGTGAG CGCTGCGGGC GCGGCGCGGG GCTTTGTGCG CTCCGCAGTG TGCGCGAGGG GAGCGCGGCC
1041 GGGGGCGGTG CCCCGCGGTG CGGGGGGGGC TGCGAGGGGA ACAAAGGCTG CGTGCGGGGT GTGTGCGTGG GGGGGTGAGC
1121 AGGGGGTGTG GGCGCGTCGG TCGGGCTGCA ACCCCCCCTG CACCCCCCTC CCCGAGTTGC TGAGCACGGC CCGGCTTCGG
1201 GTGCGGGGCT CCGTACGGGG CGTGGCGCGG GGCTCGCCGT GCCGGGCGGG GGGTGGCGGC AGGTGGGGGT GCCGGGCGGG
1281 GCGGGGCCGC CTCGGGCCGG GGAGGGCTCG GGGGAGGGGC GCGGCGGCCC CCGGAGCGCC GGCGGCTGTC GAGGCGCGGC
1361 GAGCCGCAGC CATTGCCTTT TATGGTAATC GTGCGAGAGG GCGCAGGGAC TTCCTTTGTC CCAAATCTGT GCGGAGCCGA
1441 AATCTGGGAG GCGCCGCCGC ACCCCCTCTA GCGGGCGCGG GGCGAAGCGG TGCGGCGCCG GCAGGAAGGA AATGGGCGGG
1521 GAGGGCCTTC GTGCGTCGCC GCGCCGCCGT CCCCTTCTCC CTCTCCAGCC TCGGGGCTGT CCGCGGGGGG ACGGCTGCCT
1601 TCGGGGGGGA CGGGGCAGGG CGGGGTTCGG CTTCTGGCGT GTGACCGGCG GCTCTAGAGC CTCTGCTAAC CATGTTCATG
1681 CCTTCTTCTT TTTCCTACAG CTCCTGGGCA ACGTGCTGGT TATTGTGCTG TCTCATCATT TTGGCAAAGA ATTGCAAGTT
1761 TGTACAAAAA AGCAGGCTGC CACCATGGTG AGCAAGGGCG AGGAGCTGTT CACCGGGGTG GTGCCCATCC TGGTCGAGCT
1841 GGACGGCGAC GTAAACGGCC ACAAGTTCAG CGTGTCCGGC GAGGGCGAGG GCGATGCCAC CTACGGCAAG CTGACCCTGA
1921 AGTTCATCTG CACCACCGGC AAGCTGCCCG TGCCCTGGCC CACCCTCGTG ACCACCCTGA CCTACGGCGT GCAGTGCTTC
2001 AGCCGCTACC CCGACCACAT GAAGCAGCAC GACTTCTTCA AGTCCGCCAT GCCCGAAGGC TACGTCCAGG AGCGCACCAT
2081 CTTCTTCAAG GACGACGGCA ACTACAAGAC CCGCGCCGAG GTGAAGTTCG AGGGCGACAC CCTGGTGAAC CGCATCGAGC
2161 TGAAGGGCAT CGACTTCAAG GAGGACGGCA ACATCCTGGG GCACAAGCTG GAGTACAACT ACAACAGCCA CAACGTCTAT
2241 ATCATGGCCG ACAAGCAGAA GAACGGCATC AAGGTGAACT TCAAGATCCG CCACAACATC GAGGACGGCA GCGTGCAGCT
2321 CGCCGACCAC TACCAGCAGA ACACCCCCAT CGGCGACGGC CCCGTGCTGC TGCCCGACAA CCACTACCTG AGCACCCAGT
2401 CCGCCCTGAG CAAAGACCCC AACGAGAAGC GCGATCACAT GGTCCTGCTG GAGTTCGTGA CCGCCGCCGG GATCACTCTC
2481 GGCATGGACG AGCTGTACAA GTAAACCCAG CTTTCTTGTA CAAAGTGGTG ATGGCCGGCC GCTTCGAGCA GACATGATAA
2561 GATACATTGA TGAGTTTGGA CAAACCACAA CTAGAATGCA GTGAAAAAAA TGCTTTATTT GTGAAATTTG TGATGCTATT
2641 GCTTTATTTG TAACCATTAT AAGCTGCAAT AAACAAGTTA ACAACAACAA TTGCATTCAT TTTATGTTTC AGGTTCAGGG
2721 GGAGGTGTGG GAGGTTTTTT AAAGCAAGTA AAACCTCTAC AAATGTGGTA GCGGCCGCGG CGCTCTTCCG CTTCCTCGCT
2801 CACTGACTCG CTGCGCTCGG TCGTTCGGCT GCGGCGAGCG GTATCAGCTC ACTCAAAGGC GGTAATACGG TTATCCACAG
2881 AATCAGGGGA TAACGCAGGA AAGAACATGT GAGCAAAAGG CCAGCAAAAG GCCAGGAACC GTAAAAAGGC CGCGTTGCTG
2961 GCGTTTTTCC ATAGGCTCCG CCCCCCTGAC GAGCATCACA AAAATCGACG CTCAAGTCAG AGGTGGCGAA ACCCGACAGG
3041 ACTATAAAGA TACCAGGCGT TTCCCCCTGG AAGCTCCCTC GTGCGCTCTC CTGTTCCGAC CCTGCCGCTT ACCGGATACC
3121 TGTCCGCCTT TCTCTCTTCG GGAAGCGTGG CGCTTTCTCA TAGCTCACGC TGTAGGTATC TCAGTTCGGT GTAGGTCGTT
3201 CGCTCCAAGC TGGGCTGTGT GCACGAACCC CCCGTTCAGC CCGACCGCTG CGCCTTATCC GGTAACTATC GTCTTGAGTC
3281 CAACCCGGTA AGACACGACT TATCGCCACT GGCAGCAGCC ACTGGTAACA GGATTAGCAG AGCGAGGTAT GTAGGCGGTG
3361 CTACAGAGTT CTTGAAGTGG TGGCCTAACT ACGGCTACAC TAGAAGAACA GTATTTGGTA TCTGCGCTCT GCTGAAGCCA
3441 GTTACCTTCG GAAAAAGAGT TGGTAGCTCT TGATCCGGCA AACAAACCAC CGCTGGTAGC GGTGGTTTTT TTGTTTGCAA
3521 GCAGCAGATT ACGCGCAGAA AAAAAGGATC TCAAGAAGAT CCTTTGATCT TTTCTACGGG GTCTGACGCT CAGTGGAACG
3601 AAAACTCACG TTAAGGGATT TTGGTCATGA GATTATCAAA AAGGATCTTC ACCTAGATCC TTTTAAATTA AAAATGAAGT
3681 TTTAAATCAA TCTAAAGTAT ATATGAGTAA ACTTGGTCTG ACAGTTACCA ATGCTTAATC AGTGAGGCAC CTATCTCAGC
3761 GATCTGTCTA TTTCGTTCAT CCATAGTTGC CTGACTCCCC GTCGTGTAGA TAACTACGAT ACGGGAGGGC TTACCATCTG
3841 GCCCCAGTGC TGCAATGATA CCGCGAGACC CACGCTCACC GGCTCCAGAT TTATCAGCAA TAAACCAGCC AGCCGGAAGG
3921 GCCGAGCGCA GAAGTGGTCC TGCAACTTTA TCCGCCTCCA TCCAGTCTAT TAATTGTTGC CGGGAAGCTA GAGTAAGTAG
4001 TTCGCCAGTT AATAGTTTGC GCAACGTTGT TGCCATTGCT ACAGGCATCG TGGTGTCACG CTCGTCGTTT GGTATGGCTT
4081 CATTCAGCTC CGGTTCCCAA CGATCAAGGC GAGTTACATG ATCCCCCATG TTGTGCAAAA AAGCGGTTAG CTCCTTCGGT
4161 CCTCCGATCG TTGTCAGAAG TAAGTTGGCC GCAGTGTTAT CACTCATGGT TATGGCAGCA CTGCATAATT CTCTTACTGT
4241 CATGCCATCC GTAAGATGCT TTTCTGTGAC TGGTGAGTAC TCAACCAAGT CATTCTGAGA ATAGTGTATG CGGCGACCGA
4321 GTTGCTCTTG CCCGGCGTCA ATACGGGATA ATACCGCGCC ACATAGCAGA ACTTTAAAAG TGCTCATCAT TGGAAAACGT
4401 TCTTCGGGGC GAAAACTCTC AAGGATCTTA CCGCTGTTGA GATCCAGTTC GATGTAACCC ACTCGTGCAC CCAACTGATC
4481 TTCAGCATCT TTTACTTTCA CCAGCGTTTC TGGGTGAGCA AAAACAGGAA GGCAAAATGC CGCAAAAAAG GGAATAAGGG
```

```
4561 CGACACGGAA ATGTTGAATA CTCATACTCT TCCTTTTTCA ATATTATTGA AGCATTTATC AGGGTTATTG TCTCATGAGC
4641 GGATACATAT TTGAATGTAT TTAGAAAAAT AAACAAATAG GGGTTCCGCG CACATTTCCC CGAAAAGTGC CACCTGACGT
4721 CTAAGAAACC ATTATTATCA TGACATTAAC CTATAAAAAT AGGCGTATCA CGAGGCCCTT TCGTCGGCGC GCCGCGGCCG
4801 C
```

### Validation by Restriction Enzyme Digestion

| Restriction Enzymes | Cutting Sites | DNA Fragments (bp) |
|---|---|---|
| **NdeI** | 275 | 4801 |
| **NaeI** | 1341, 1500, 2537 | 159, 1037, 3605 |
| **HpaI** | 2680 | 4801 |
| **SpeI** | 40 | 4801 |
| **ApaLI** | 3220, 4466 | 1246, 3555 |
| **ApaLI+NdeI** | 275, 3220, 4466 | 2945, 1246, 610 |
| **ApaLI+SpeI** | 40, 3220, 4466 | 3180, 1246, 375 |
| **ApaLI+NaeI** | 1341, 1500, 2537, 3220, 4466 | 159, 1037, 683, 1246, 1676 |
| **ApaLI+HpaI** | 2680, 3220, 4466 | 540, 1246, 3015 |

### Vector Summary

| | |
|---|---|
| Vector ID | VB220302-1045eru |
| Vector Name | pRP[shRNA]-EGFP-U6>{hsa-microRNA-502-3p} |
| Vector Size | 3861 bp |
| Vector Type | Mammalian shRNA knockdown vector |
| Inserted shRNA | {hsa-microRNA-502-3p} |
| Target Sequence | TGAATCCTTGCCCAGGTGCATT |
| Inserted Marker | EGFP |
| Plasmid Copy Number | High |
| Antibiotic Resistance | Ampicillin |
| Cloning Host | VB UltraStable (or alternative strain) |

### Vector Map

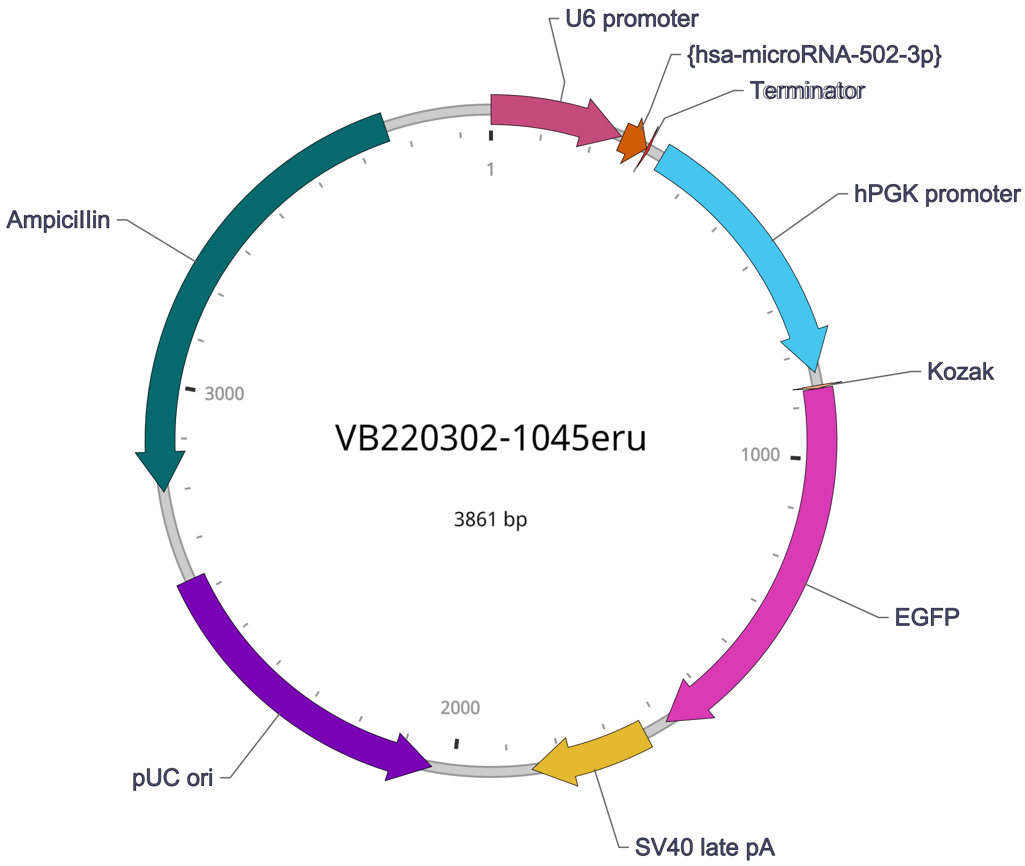


### Vector Components

| Name | Position | Size (bp) | Type | Description | Application notes |
|---|---|---|---|---|---|
| U6 promoter | 1-249 | 249 | Promoter | Human U6 small nuclear 1 promoter | Pol III promoter; drives expression of small RNAs. |
| **{hsa-microRNA-502-3p}** | 252-301 | 50 | shRNA | *None* | *None* |
| Terminator | 302-306 | 5 | terminator | Pol III transcription terminator | Allows transcription termination of small RNA transcribed by Pol III RNA polymerase. |
| hPGK promoter | 334-838 | 505 | Promoter | Human phosphoglycerate kinase 1 promoter | Medium-strength promoter. |
| Kozak | 863-868 | 6 | Miscellaneous | Kozak translation initiation sequence | Facilitates translation initiation of ATG start codon downstream of the Kozak sequence. |
| **EGFP** | 869-1588 | 720 | CDS | Enhanced green fluorescent protein; codon optimized based on a variant of wild type GFP from the jellyfish Aequorea victoria | Commonly used green fluorescent protein; ranked high in brightness, photostability and pH stability among all fluorescent proteins. |
| SV40 late pA | 1633-1854 | 222 | PolyA_signal | Simian virus 40 late polyadenylation signal | Allows transcription termination and polyadenylation of mRNA transcribed by Pol II RNA polymerase. |
| pUC ori | complement (2042-2630) | 589 | Rep_origin | pUC origin of replication | Facilitates plasmid replication in E. coli; regulates high-copy plasmid number (500-700). |
| AmpicilIin | complement (2801-3661) | 861 | CDS | AmpicilIin resistance gene | Allows E. coli to be resistant to ampicilIin. |

**Note:** Components added by user are listed in **bold red** text.

### Vector Sequence

1

81 GAGGGCCTAT TTCCCATGAT TCCTTCATAT TTGCATATAC GATACAAGGC TGTTAGAGAG ATAATTGGAA TTAATTTGAC

161 TGTAAACACA AAGATATTAG TACAAAATAC GTGACGTAGA AAGTAATAAT TTCTTGGGTA GTTTGCAGTT TTAAAATTAT

241 GTTTTAAAAT GGACTATCAT ATGCTTACCG TAACTTGAAA GTATTTCGAT TTCTTGGCTT TATATATCTT GTGGAAAGGA

321 CGAAACACCG GTGAATCCTT GCCCAGGTGC ATTCTCGAGA ATGCACCTGG GCAAGGATTC ATTTTTGAAT TCCAACTTTG

401 TATAGAAAAG TTGGGGTTGC GCCTTTTCCA AGGCAGCCCT GGGTTTGCGC AGGGACGCGG CTGCTCTGGG CGTGGTTCCG

481 GGAAACGCAG CGGCGCCGAC CCTGGGTCTC GCACATTCTT CACGTCCGTT CGCAGCGTCA CCCGGATCTT CGCCGCTACC

561 CTTGTGGGCC CCCCGGCGAC GCTTCCTGCT CCGCCCCTAA GTCGGGAAGG TTCCTTGCGG TTCGCGGCGT GCCGGACGTG

641 ACAAACGGAA GCCGCACGTC TCACTAGTAC CCTCGCAGAC GGACAGCGCC AGGGAGCAAT GGCAGCGCGC CGACCGCGAT

721 GGGCTGTGGC CAATAGCGGC TGCTCAGCAG GGCGCGCCGA GAGCAGCGGC CGGGAAGGGG CGGTGCGGGA GGCGGGGTGT

801 GGGGCGGTAG TGTGGGCCCT GTTCCTGCCC GCGCGGTGTT CCGCATTCTG CAAGCCTCCG GAGCGCACGT CGGCAGTCGG

881 CTCCCTCGTT GACCGAATCA CCGACCTCTC TCCCCAGGCA AGTTTGTACA AAAAAGCAGG CTGCCACCAT GGTGAGCAAG

961 GGCGAGGAGC TGTTCACCGG GGTGGTGCCC ATCCTGGTCG AGCTGGACGG CGACGTAAAC GGCCACAAGT TCAGCGTGTC

1041 CGGCGAGGGC GAGGGCGATG CCACCTACGG CAAGCTGACC CTGAAGTTCA TCTGCACCAC CGGCAAGCTG CCCGTGCCCT

1121 GGCCCACCCT CGTGACCACC CTGACCTACG GCGTGCAGTG CTTCAGCCGC TACCCCGACC ACATGAAGCA GCACGACTTC

1201 TTCAAGTCCG CCATGCCCGA AGGCTACGTC CAGGAGCGCA CCATCTTCTT CAAGGACGAC GGCAACTACA AGACCCGCGC

1281 CGAGGTGAAG TTCGAGGGCG ACACCCTGGT GAACCGCATC GAGCTGAAGG GCATCGACTT CAAGGAGGAC GGCAACATCC

1361 TGGGGCACAA GCTGGAGTAC AACTACAACA GCCACAACGT CTATATCATG GCCGACAAGC AGAAGAACGG CATCAAGGTG

1441 AACTTCAAGA TCCGCCACAA CATCGAGGAC GGCAGCGTGC AGCTCGCCGA CCACTACCAG CAGAACACCC CCATCGGCGA

1521 CGGCCCCGTG CTGCTGCCCG ACAACCACTA CCTGAGCACC CAGTCCGCCC TGAGCAAAGA CCCCAACGAG AAGCGCGATC

1601 ACATGGTCCT GCTGGAGTTC GTGACCGCCG CCGGGATCAC TCTCGGCATG GACGAGCTGT ACAAGTAAAC CCAGCTTTCT

1681 TGTACAAAGT GGTGATGGCC GGCCGCTTCG AGCAGACATG ATAAGATACA TTGATGAGTT TGGACAAACC ACAACTAGAA

1761 TGCAGTGAAA AAAATGCTTT ATTTGTGAAA TTTGTGATGC TATTGCTTTA TTTGTAACCA TTATAAGCTG CAATAAACAA

1841 GTTAACAACA ACAATTGCAT TCATTTTATG TTTCAGGTTC AGGGGGAGGT GTGGGAGGTT TTTTAAAGCA AGTAAAACCT

1921 CTACAAATGT GGTAGGCGCT CTTCCGCTTC CTCGCTCACT GACTCGCTGC GCTCGGTCGT TCGGCTGCGG CGAGCGGTAT

2001 CAGCTCACTC AAAGGCGGTA ATACGGTTAT CCACAGAATC AGGGGATAAC GCAGGAAAGA ACATGTGAGC AAAAGGCCAG

2081 CAAAAGGCCA GGAACCGTAA AAAGGCCGCG TTGCTGGCGT TTTTCCATAG GCTCCGCCCC CCTGACGAGC ATCACAAAAA

2161 TCGACGCTCA AGTCAGAGGT GGCGAAACCC GACAGGACTA TAAAGATACC AGGCGTTTCC CCCTGGAAGC TCCCTCGTGC

2241 GCTCTCCTGT TCCGACCCTG CCGCTTACCG GATACCTGTC CGCCTTTCTC CCTTCGGGAA GCGTGGCGCT TTCTCATAGC

2321 TCACGCTGTA GGTATCTCAG TTCGGTGTAG GTCGTTCGCT CCAAGCTGGG CTGTGTGCAC GAACCCCCCG TTCAGCCCGA

2401 CCGCTGCGCC TTATCCGGTA ACTATCGTCT TGAGTCCAAC CCGGTAAGAC ACGACTTATC GCCACTGGCA GCAGCCACTG

2481 GTAACAGGAT TAGCAGAGCG AGGTATGTAG GCGGTGCTAC AGAGTTCTTG AAGTGGTGGC CTAACTACGG CTACACTAGA

2561 AGAACAGTAT TTGGTATCTG CGCTCTGCTG AAGCCAGTTA CCTTCGGAAA AAGAGTTGGT AGCTCTTGAT CCGGCAAACA

2641 AACCACCGCT GGTAGCGGTG GTTTTTTTGT TTGCAAGCAG CAGATTACGC GCAGAAAAAA AGGATCTCAA GAAGATCCTT

2721 TGATCTTTTC TACGGGGTCT GACGCTCAGT GGAACGAAAA CTCACGTTAA GGGATTTTGG TCATGAGATT ATCAAAAAGG

2801 ATCTTCACCT AGATCCTTTT AAATTAAAAA TGAAGTTTTA AATCAATCTA AAGTATATAT GAGTAAACTT GGTCTGACAG

2881 TTACCAATGC TTAATCAGTG AGGCACCTAT CTCAGCGATC TGTCTATTTC GTTCATCCAT AGTTGCCTGA CTCCCCGTCG

2961 TGTAGATAAC TACGATACGG GAGGGCTTAC CATCTGGCCC CAGTGCTGCA ATGATACCGC GAGATCCACG CTCACCGGCT

3041 CCAGATTTAT CAGCAATAAA CCAGCCAGCC GGAAGGGCCG AGCGCAGAAG TGGTCCTGCA ACTTTATCCG CCTCCATCCA

3121 GTCTATTAAT TGTTGCCGGG AAGCTAGAGT AAGTAGTTCG CCAGTTAATA GTTTGCGCAA CGTTGTTGCC ATTGCTACAG

3201 GCATCGTGGT GTCACGCTCG TCGTTTGGTA TGGCTTCATT CAGCTCCGGT TCCCAACGAT CAAGGCGAGT TACATGATCC

3281 CCCATGTTGT GCAAAAAAGC GGTTAGCTCC TTCGGTCCTC CGATCGTTGT CAGAAGTAAG TTGGCCGCAG TGTTATCACT

3361 CATGGTTATG GCAGCACTGC ATAATTCTCT TACTGTCATG CCATCCGTAA GATGCTTTTC TGTGACTGGT GAGTACTCAA

3441 CCAAGTCATT CTGAGAATAG TGTATGCGGC GACCGAGTTG CTCTTGCCCG GCGTCAATAC GGGATAATAC CGCGCCACAT

3521 AGCAGAACTT TAAAAGTGCT CATCATTGGA AAACGTTCTT CGGGGCGAAA ACTCTCAAGG ATCTTACCGC TGTTGAGATC

3601 CAGTTCGATG TAACCCACTC GTGCACCCAA CTGATCTTCA GCATCTTTTA CTTTCACCAG CGTTTCTGGG TGAGCAAAAA

3681 CAGGAAGGCA AAATGCCGCA AAAAAGGGAA TAAGGGCGAC ACGGAAATGT TGAATACTCA TACTCTTCCT TTTTCAATAT

3761 TATTGAAGCA TTTATCAGGG TTATTGTCTC ATGAGCGGAT ACATATTTGA ATGTATTTAG AAAAATAAAC AAATAGGGGT

TCCGCGCACA TTTCCCCGAA AAGTGCCACC TGACGTCTAA GAAACCATTA TTATCATGAC ATTAACCTAT AAAAATAGGC

3841 GTATCACGAG GCCCTTTCGT C

### Vector Summary

| | |
|---|---|
| Vector ID | VB220311-1069att |
| Vector Name | pRP[Exp]-CAG>EGFP:miRNA-502-3p sponge |
| Vector Size | 5041 bp |
| Vector Type | Mammalian Gene Expression Vector |
| Plasmid Copy Number | High |
| Antibiotic Resistance | Ampicillin |
| Cloning Host | VB UltraStable (or alternative strain) |

### Vector Map

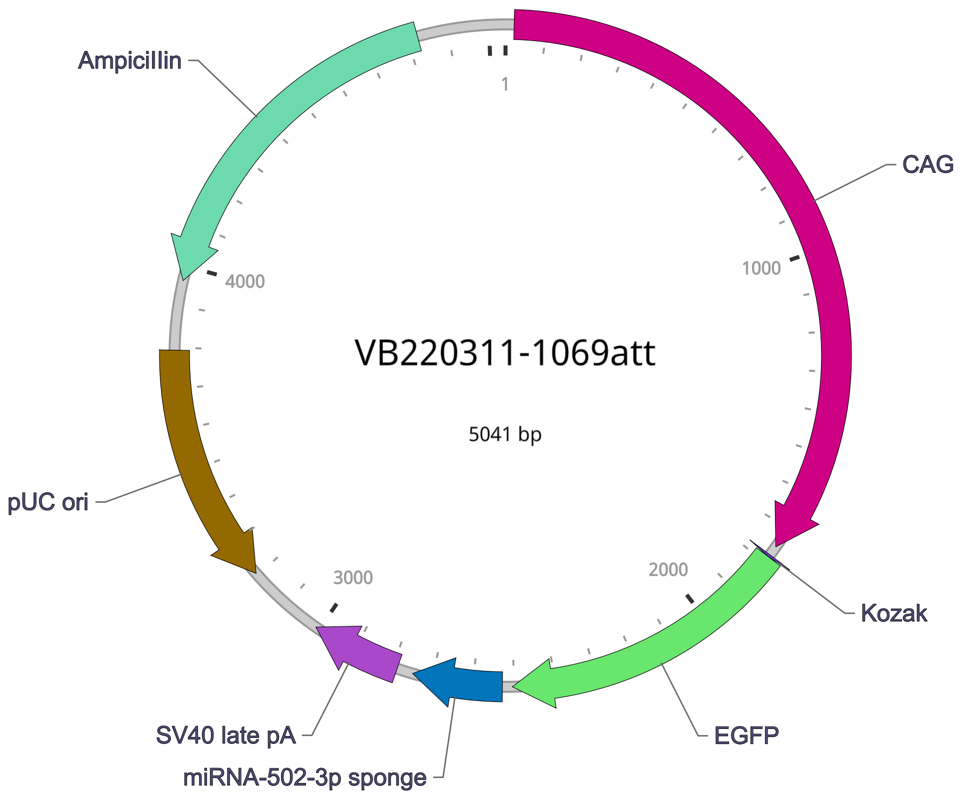


### Vector Components

| Name | Position | Size (bp) | Type | Description | Application notes |
|---|---|---|---|---|---|

| Name | Position | Size (bp) | Type | Description | Application notes |
|---|---|---|---|---|---|
| CAG | 22-1754 | 1733 | Promoter | CMV early enhancer fused to modified chicken beta-actin promoter | Strong promoter. |
| Kozak | 1779-1784 | 6 | Miscellaneous | Kozak translation initiation sequence | Facilitates translation initiation of ATG start codon downstream of the Kozak sequence. |
| EGFP | 1785-2504 | 720 | CDS | Enhanced green fluorescent protein; codon optimized based on a variant of wild type GFP from the jellyfish Aequorea victoria | Commonly used green fluorescent protein; ranked high in brightness, photostability and pH stability among all fluorescent proteins. |
| miRNA-502-3p sponge | 2529-2749 | 221 | misc_feature | *None* | *None* |
| SV40 late pA | 2789-3010 | 222 | PolyA_signal | Simian virus 40 late polyadenylation signal | Allows transcription termination and polyadenylation of mRNA transcribed by Pol II RNA polymerase. |
| pUC ori | complement (3206-3794) | 589 | Rep_origin | pUC origin of replication | Facilitates plasmid replication in E. coli; regulates high-copy plasmid number (500-700). |
| AmpicilIin | complement (3965-4825) | 861 | CDS | AmpicilIin resistance gene | Allows E. coli to be resistant to ampicillin. |

**Note:** Components added by user are listed in **bold red** text.

### Vector Sequence

```
  1 CAACTTTGTA TAGAAAAGTT GCTCGACATT GATTATTGAC TAGTTATTAA TAGTAATCAA TTACGGGGTC ATTAGTTCAT
 81 AGCCCATATA TGGAGTTCCG CGTTACATAA CTTACGGTAA ATGGCCCGCC TGGCTGACCG CCCAACGACC CCCGCCCATT
161 GACGTCAATA ATGACGTATG TTCCCATAGT AACGCCAATA GGGACTTTCC ATTGACGTCA ATGGGTGGAG TATTTACGGT
241 AAACTGCCCA CTTGGCAGTA CATCAAGTGT ATCATATGCC AAGTACGCCC CCTATTGACG TCAATGACGG TAAATGGCCC
321 GCCTGGCATT ATGCCCAGTA CATGACCTTA TGGGACTTTC CTACTTGGCA GTACATCTAC GTATTAGTCA TCGCTATTAC
401 CATGGTCGAG GTGAGCCCCA CGTTCTGCTT CACTCTCCCC ATCTCCCCCC CCTCCCCACC CCCAATTTTG TATTTATTTA
481 TTTTTTAATT ATTTTGTGCA GCGATGGGGG CGGGGGGGGG GGGGGGGCGC GCGCCAGGCG GGGCGGGGCG GGGCGAGGGG
```

561 CGGGGCGGGG CGAGGCGGAG AGGTGCGGCG GCAGCCAATC AGAGCGGCGC GCTCCGAAAG TTTCCTTTTA TGGCGAGGCG

641 GCGGCGGCGG CGGCCCTATA AAAAGCGAAG CGCGCGGCGG GCGGGAGTCG CTGCGCGCTG CCTTCGCCCC GTGCCCCGCT

721 CCGCCGCCGC CTCGCGCCGC CCGCCCCGGC TCTGACTGAC CGCGTTACTC CCACAGGTGA GCGGGCGGGA CGGCCCTTCT

801 CCTCCGGGCT GTAATTAGCG CTTGGTTTAA TGACGGCTTG TTTCTTTTCT GTGGCTGCGT GAAAGCCTTG AGGGGCTCCG

881 GGAGGGCCCT TTGTGCGGGG GGAGCGGCTC GGGGGGTGCG TGCGTGTGTG TGTGCGTGGG GAGCGCCGCG TGCGGCTCCG

961 CGCTGCCCGG CGGCTGTGAG CGCTGCGGGC GCGGCGCGGG GCTTTGTGCG CTCCGCAGTG TGCGCGAGGG GAGCGCGGCC

1041 GGGGGCGGTG CCCCGCGGTG CGGGGGGGGC TGCGAGGGGA ACAAAGGCTG CGTGCGGGGT GTGTGCGTGG GGGGGTGAGC

1121 AGGGGGTGTG GGCGCGTCGG TCGGGCTGCA ACCCCCCCTG CACCCCCCTC CCCGAGTTGC TGAGCACGGC CCGGCTTCGG

1201 GTGCGGGGCT CCGTACGGGG CGTGGCGCGG GGCTCGCCGT GCCGGGCGGG GGGTGGCGGC AGGTGGGGGT GCCGGGCGGG

1281 GCGGGGCCGC CTCGGGCCGG GGAGGGCTCG GGGGAGGGGC GCGGCGGCCC CCGGAGCGCC GGCGGCTGTC GAGGCGCGGC

1361 GAGCCGCAGC CATTGCCTTT TATGGTAATC GTGCGAGAGG GCGCAGGGAC TTCCTTTGTC CCAAATCTGT GCGGAGCCGA

1441 AATCTGGGAG GCGCCGCCGC ACCCCCTCTA GCGGGCGCGG GGCGAAGCGG TGCGGCGCCG GCAGGAAGGA AATGGGCGGG

1521 GAGGGCCTTC GTGCGTCGCC GCGCCGCCGT CCCCTTCTCC CTCTCCAGCC TCGGGGCTGT CCGCGGGGGG ACGGCTGCCT

1601 TCGGGGGGGA CGGGGCAGGG CGGGGTTCGG CTTCTGGCGT GTGACCGGCG GCTCTAGAGC CTCTGCTAAC CATGTTCATG

1681 CCTTCTTCTT TTTCCTACAG CTCCTGGGCA ACGTGCTGGT TATTGTGCTG TCTCATCATT TTGGCAAAGA ATTGCAAGTT

1761 TGTACAAAAA AGCAGGCTGC CACCATGGTG AGCAAGGGCG AGGAGCTGTT CACCGGGGTG GTGCCCATCC TGGTCGAGCT

1841 GGACGGCGAC GTAAACGGCC ACAAGTTCAG CGTGTCCGGC GAGGGCGAGG GCGATGCCAC CTACGGCAAG CTGACCCTGA

1921 AGTTCATCTG CACCACCGGC AAGCTGCCCG TGCCCTGGCC CACCCTCGTG ACCACCCTGA CCTACGGCGT GCAGTGCTTC

2001 AGCCGCTACC CCGACCACAT GAAGCAGCAC GACTTCTTCA AGTCCGCCAT GCCCGAAGGC TACGTCCAGG AGCGCACCAT

2081 CTTCTTCAAG GACGACGGCA ACTACAAGAC CCGCGCCGAG GTGAAGTTCG AGGGCGACAC CCTGGTGAAC CGCATCGAGC

2161 TGAAGGGCAT CGACTTCAAG GAGGACGGCA ACATCCTGGG GCACAAGCTG GAGTACAACT ACAACAGCCA CAACGTCTAT

2241 ATCATGGCCG ACAAGCAGAA GAACGGCATC AAGGTGAACT TCAAGATCCG CCACAACATC GAGGACGGCA GCGTGCAGCT

2321 CGCCGACCAC TACCAGCAGA ACACCCCCAT CGGCGACGGC CCCGTGCTGC TGCCCGACAA CCACTACCTG AGCACCCAGT

2401 CCGCCCTGAG CAAAGACCCC AACGAGAAGC GCGATCACAT GGTCCTGCTG GAGTTCGTGA CCGCCGCCGG GATCACTCTC

2481 GGCATGGACG AGCTGTACAA GTAAACCCAG CTTTCTTGTA CAAAGTGGCT CGAGCCGGTG AATCCTTGGG TGGTGCATTC

2561 CGGTGAATCC TTGGGTGGTG CATTCCGGTG AATCCTTGGG TGGTGCATTC CGGTGAATCC TTGGGTGGTG CATTCCGGTG

2641 AATCCTTGGG TGGTGCATTC CGGTGAATCC TTGGGTGGTG CATTCCGGTG AATCCTTGGG TGGTGCATTC CGGTGAATCC

2721 TTGGGTGGTG CATTCCGGAT CGCGGGCCCC AACTTTATTA TACATAGTTG ATGGCCGGCC GCTTCGAGCA GACATGATAA

2801 GATACATTGA TGAGTTTGGA CAAACCACAA CTAGAATGCA GTGAAAAAAA TGCTTTATTT GTGAAATTTG TGATGCTATT

2881 GCTTTATTTG TAACCATTAT AAGCTGCAAT AAACAAGTTA ACAACAACAA TTGCATTCAT TTTATGTTTC AGGTTCAGGG

2961 GGAGGTGTGG GAGGTTTTTT AAAGCAAGTA AAACCTCTAC AAATGTGGTA GCGGCCGCGG CGCTCTTCCG CTTCCTCGCT

3041 CACTGACTCG CTGCGCTCGG TCGTTCGGCT GCGGCGAGCG GTATCAGCTC ACTCAAAGGC GGTAATACGG TTATCCACAG

3121 AATCAGGGGA TAACGCAGGA AAGAACATGT GAGCAAAAGG CCAGCAAAAG GCCAGGAACC GTAAAAAGGC CGCGTTGCTG

3201 GCGTTTTTCC ATAGGCTCCG CCCCCCTGAC GAGCATCACA AAAATCGACG CTCAAGTCAG AGGTGGCGAA ACCCGACAGG

3281 ACTATAAAGA TACCAGGCGT TTCCCCCTGG AAGCTCCCTC GTGCGCTCTC CTGTTCCGAC CCTGCCGCTT ACCGGATACC

3361 TGTCCGCCTT TCTCTCTTCG GGAAGCGTGG CGCTTTCTCA TAGCTCACGC TGTAGGTATC TCAGTTCGGT GTAGGTCGTT

3441 CGCTCCAAGC TGGGCTGTGT GCACGAACCC CCCGTTCAGC CCGACCGCTG CGCCTTATCC GGTAACTATC GTCTTGAGTC

3521 CAACCCGGTA AGACACGACT TATCGCCACT GGCAGCAGCC ACTGGTAACA GGATTAGCAG AGCGAGGTAT GTAGGCGGTG

3601 CTACAGAGTT CTTGAAGTGG TGGCCTAACT ACGGCTACAC TAGAAGAACA GTATTTGGTA TCTGCGCTCT GCTGAAGCCA

3681 GTTACCTTCG GAAAAAGAGT TGGTAGCTCT TGATCCGGCA AACAAACCAC CGCTGGTAGC GGTGGTTTTT TTGTTTGCAA

3761 GCAGCAGATT ACGCGCAGAA AAAAAGGATC TCAAGAAGAT CCTTTGATCT TTTCTACGGG GTCTGACGCT CAGTGGAACG

3841 AAAACTCACG TTAAGGGATT TTGGTCATGA GATTATCAAA AAGGATCTTC ACCTAGATCC TTTTAAATTA AAAATGAAGT

3921 TTTAAATCAA TCTAAAGTAT ATATGAGTAA ACTTGGTCTG ACAGTTACCA ATGCTTAATC AGTGAGGCAC CTATCTCAGC

4001 GATCTGTCTA TTTCGTTCAT CCATAGTTGC CTGACTCCCC GTCGTGTAGA TAACTACGAT ACGGGAGGGC TTACCATCTG

4081 GCCCCAGTGC TGCAATGATA CCGCGAGACC CACGCTCACC GGCTCCAGAT TTATCAGCAA TAAACCAGCC AGCCGGAAGG

4161 GCCGAGCGCA GAAGTGGTCC TGCAACTTTA TCCGCCTCCA TCCAGTCTAT TAATTGTTGC CGGGAAGCTA GAGTAAGTAG

4241 TTCGCCAGTT AATAGTTTGC GCAACGTTGT TGCCATTGCT ACAGGCATCG TGGTGTCACG CTCGTCGTTT GGTATGGCTT

4321 CATTCAGCTC CGGTTCCCAA CGATCAAGGC GAGTTACATG ATCCCCCATG TTGTGCAAAA AAGCGGTTAG CTCCTTCGGT

```
4401 CCTCCGATCG TTGTCAGAAG TAAGTTGGCC GCAGTGTTAT CACTCATGGT TATGGCAGCA CTGCATAATT CTCTTACTGT
4481 CATGCCATCC GTAAGATGCT TTTCTGTGAC TGGTGAGTAC TCAACCAAGT CATTCTGAGA ATAGTGTATG CGGCGACCGA
4561 GTTGCTCTTG CCCGGCGTCA ATACGGGATA ATACCGCGCC ACATAGCAGA ACTTTAAAAG TGCTCATCAT TGGAAAACGT
4641 TCTTCGGGGC GAAAACTCTC AAGGATCTTA CCGCTGTTGA GATCCAGTTC GATGTAACCC ACTCGTGCAC CCAACTGATC
4721 TTCAGCATCT TTTACTTTCA CCAGCGTTTC TGGGTGAGCA AAAACAGGAA GGCAAAATGC CGCAAAAAAG GGAATAAGGG
4801 CGACACGGAA ATGTTGAATA CTCATACTCT TCCTTTTTCA ATATTATTGA AGCATTTATC AGGGTTATTG TCTCATGAGC
4881 GGATACATAT TTGAATGTAT TTAGAAAAAT AAACAAATAG GGGTTCCGCG CACATTTCCC CGAAAAGTGC CACCTGACGT
4961 CTAAGAAACC ATTATTATCA TGACATTAAC CTATAAAAAT AGGCGTATCA CGAGGCCCTT TCGTCGGCGC GCCGCGGCCG
5041 C
```

### Validation by Restriction Enzyme Digestion

| Restriction Enzymes | Cutting Sites | DNA Fragments (bp) |
|---|---|---|
| **XhoI** | 2530 | 5041 |
| **NdeI** | 275 | 5041 |
| **HpaI** | 2920 | 5041 |
| **SpeI** | 40 | 5041 |
| **ApaLI** | 3460, 4706 | 1246, 3795 |
| **ApaLI+XhoI** | 2530, 3460, 4706 | 930, 1246, 2865 |
| **ApaLI+NdeI** | 275, 3460, 4706 | 3185, 1246, 610 |
| **ApaLI+SpeI** | 40, 3460, 4706 | 3420, 1246, 375 |
| **ApaLI+HpaI** | 2920, 3460, 4706 | 540, 1246, 3255 |
