## Supplementary material for "MicroRNA-502-3p modulates the GABA A subunits, synaptic proteins and mitochondrial morphology in hippocampal neurons": SI File 2

**Supplementary Information**

**Table 1. Oligonucleotide sequences of primers used for quantitative reverse transcription-polymerase chain reaction analysis**

| **Gene(s)** | **Sequence(s)** |
| --- | --- |
| GABRA1 | F- 5^'^ TGTGTGAAAGTGTTTAAAAGGGCA 3^'^ |
|  | R- 5^'^ TCAGCTTCAACATTGACGAATAAA 3^'^ |
| U6 SnRNA | F- 5^'^ CGCTTCGGCAGCACATATACTAA 3^'^ |
|  | R- 5^'^ TATGGAACGCTTCACGAATTTGC 3^'^ |
| miR-502-3p | F- 5^'^ AATGCACCTGGGCAAGGATTCA 3^'^ |

**Table 2. Summary of antibody dilutions and conditions used in the immunostaining analysis**

| **Marker(s)** | **Primary Antibody and Dilution**  **(4^°^C, overnight)** | **Purchased from Company, City & State** | **Secondary Antibody**  **(Room temperature, 1 h)** | **Purchased from Company, City & State** |
| --- | --- | --- | --- | --- |
| GABRα1  (BS-1232R) | Rabbit polyclonal  1:200 | Bioss Antibodies,  Woburn, MA | Goat anti-rabbit IgG HRP 1:500  (A9169-2 mL) | Millipore Sigma  Burlington, MA |

**Table 3. Summary of antibody dilutions and conditions used in the immunoblotting analysis**

| **Marker(s)** | **Primary Antibody and Dilution(s)**  **(4^°^C, overnight)** | **Purchased from Company, City & State** | **Secondary Antibody, Dilution(s)**  **(Room temperature, 2 h)** | **Purchased from Company, City & State** |
| --- | --- | --- | --- | --- |
| GABRα1  (BS-1232R) | Rabbit polyclonal  1:2000 | Bioss Antibodies,  Woburn, MA | Goat anti-rabbit IgG HRP 1:10,000  (A9169-2 mL) | Millipore Sigma  Burlington, MA |
| GABRβ3  (NB300-199) | Rabbit polyclonal  1:1000 | Novus, Minneapolis, MN | Goat anti-rabbit IgG HRP 1:10,000  (A9169-2 mL) | Millipore Sigma  Burlington, MA |
| GABRγ2  (14104-1-AP) | Rabbit polyclonal  1:2000 | Proteintech, Rosemont, IL | Goat anti-rabbit IgG HRP 1:10,000  (A9169-2 mL) | Millipore Sigma  Burlington, MA |
| Calmodulin  (10541-1-AP) | Rabbit polyclonal  1:500 | Proteintech, Rosemont, IL | Goat anti-rabbit IgG HRP 1:10,000  (A9169-2 mL) | Millipore Sigma  Burlington, MA |
| SNAP25  (PA5-85396) | Rabbit polyclonal  1:1000 | Invitrogen, Waltham, MA | Goat anti-rabbit IgG HRP 1:10,000  (A9169-2 mL) | Millipore Sigma  Burlington, MA |
| Syntaxin1 (66437-1-Ig) | Mouse monoclonal 1:1000 | Proteintech, Rosemont, IL | Rabbit anti-mouse IgG HRP 1:10,000  (A9044-2 mL) | Millipore Sigma  Burlington, MA |
| MAP2 (17490-1-AP) | Rabbit polyclonal  1:1000 | Proteintech, Rosemont, IL | Goat anti-rabbit IgG HRP 1:10,000  (A9169-2 mL) | Millipore Sigma  Burlington, MA |
| NRXN1 (NBP1-00219) | Rabbit polyclonal 1:1000 | Novus, Minneapolis, MN | Goat anti-rabbit IgG HRP 1:10,000  (A9169-2 mL) | Millipore Sigma  Burlington, MA |
| VAMP2 (10135-1-AP) | Rabbit polyclonal  1:2000 | Proteintech, Rosemont, IL | Goat anti-rabbit IgG HRP 1:10,000  (A9169-2 mL) | Millipore Sigma  Burlington, MA |
| β-actin  (803001) | Mouse monoclonal 1:5000 | Millipore Sigma  Burlington, MA | Rabbit anti-mouse IgG HRP 1:10,000  (A9044-2 mL) | Millipore Sigma  Burlington, MA |
